# Modelling sex-specific whole-body metabolic responses to feeding and fasting

**DOI:** 10.1101/2024.04.29.591595

**Authors:** Stéphanie M. C. Abo, Anita T. Layton

## Abstract

Men exhibit a preference for carbohydrate metabolism, whereas women tend to favor lipid metabolism. Significant sex-based differences in energy oxidation are evident across various metabolic states, including fasting and feeding. While some of these differences can be attributed to variations in body composition—such as increased fat mass in women and higher muscle mass in men—there are also inherent disparities in metabolic fluxes. For instance, women exhibit increased rates of lipolysis independent of body composition. However, there remain gaps in our understanding of how sex influences the metabolism of specific organs and how these differences manifest at the systemic level. To address some of these gaps, we developed a sex-specific, whole-body, multi-scale model of metabolism during feeding and fasting. Our model represents healthy young adults (male and female) and integrates cellular metabolism in organs with whole-body responses following various mixed meals, particularly high-carbohydrate and high-fat meals. We explored sex-related variations in metabolic responses during both the absorptive and postabsorptive phases following meals. Our model predicted that sex-related metabolic differences observed at the systemic level are driven by variations in nutrient storage and oxidation patterns in the liver, skeletal muscle, and adipose tissue. We hypothesized that sex differences in hepatic glucose output during short-term fasts are partly influenced by variations in free fatty acids, glycerol, and glycogen handling. We also identified a candidate mechanism, possibly more prevalent in the female liver, where lipids are redirected toward carbohydrate metabolism to support hepatic glucose production. Integrating sex-specific data and parameters into multi-scale frameworks holds promise for enhancing our understanding of human metabolism and its modulation by sex.

**Author summary:** Men and women exhibit different metabolic preferences, with men favoring carbohydrate metabolism and women favoring lipid metabolism. These differences impact energy usage during fasting and feeding, influenced by body composition and inherent metabolic variations. However, there remain gaps in our understanding of how sex influences the metabolism of specific organs and how these differences manifest at the systemic level. To address these gaps, we developed a sex-specific whole-body metabolic model, representing healthy young adults. Our model explores how men and women metabolize mixed meals, especially high-carbohydrate and high-fat meals. We found that sex-related metabolic variations arise from differences in nutrient storage and utilization in key organs like the liver, muscles, and fat tissue. We propose that variations in free fatty acids and glycogen handling contribute to differences in hepatic glucose output between men and women during short-term fasting. Integrating sex-specific data into metabolic models can enhance our understanding of human metabolism and its modulation by sex.

## 1 Introduction

Obesity and related metabolic disorders, such as type 2 diabetes, have emerged as worldwide epidemics [1, 2]. Nutrition has been identified as the primary modifiable factor that can be addressed to mitigate the escalating prevalence of obesity and metabolic diseases. Nutritional consumption of humans composes largely of carbohydrates, fat, and proteins. A key factor in determining the ideal nutritional intake is the relative efficiency of substrate oxidation and conversion. Carbohydrates are preferentially oxidized over fats, which helps stabilize blood glucose levels. While carbohydrate consumption acutely boosts carbohydrate oxidation, it usually only minimally increases *de novo* lipogenesis. In humans, the storage capacity for carbohydrates is limited, but that for adipose tissue is much more extensive; as a result, fat storage is favored when there is excessive caloric intake. The high energy density of fats, providing more than twice the energy per gram compared to carbohydrates or protein, may further contribute to weight gain if not offset by increased energy expenditure.

One significant aspect influencing whole-body metabolism is the role of sex. The impact of sex on metabolic processes is a burgeoning field of research, and recent experimental evidence underscores its importance [3–6]. Sexual dimorphism is observed not only in body composition but also in metabolic rates, substrate utilization, and hormonal regulation [4, 7–13]. Sex differences in metabolism manifest across various physiological conditions: fasting [14, 15], feeding [8, 16, 17], hypoglycemia [6, 18, 19], exercise [3, 20, 21], and more [22–25]. While both sexes exhibit general responses such as hepatic glycogenolysis, gluconeogenesis, and adipose tissue lipolysis, quantitative distinctions in substrate utilization are notable [22–24]. Studies have revealed a propensity for carbohydrate metabolism in men and lipid metabolism in women, highlighting significant sex-based differences in fuel oxidation during different metabolic states [16, 26–28]. The distribution of adipose storage exhibits distinct sexual dimorphism, with men tending to accumulate more fat in the android region and women favoring the gynoid region [4]. Apart from adipose tissue, lipid storage occurs in various tissues, including the liver and skeletal muscle, exhibiting sex-specific patterns. Women tend to demonstrate greater lipid storage in muscle [3, 8], while men display more pronounced storage in the liver [17]. These sex-based disparities influence resting, postprandial, fasting, and exercise metabolism, contributing to differences in substrate utilization and the risk of metabolic diseases [4, 20]. Understanding the role of sex in mediating metabolic responses can inform tailored nutritional guidelines and therapeutic strategies, potentially enhancing glycemic control and mitigating the risk of metabolic disorders in both sexes. However, despite the acknowledged significance of sex-specific differences, integrating these nuances into comprehensive models has been a challenging yet necessary endeavor.

Many mathematical models address metabolism, spanning various scales and facets crucial for studying energy metabolism and whole-body homeostatic balance [14, 20, 21, 29–38]. For instance, there exist well developed compartment models of glucose homeostasis. Cobelli et al. [39] developed the artificial pancreas for type 1 diabetes mellitus—a simulator model of the glucose-insulin system. Approved by the Food and Drug Administration in 2013, this model can be used as a substitute for preclinical trials for select insulin treatments. An improved iteration, transitioning from simulating a single meal to a full day, was introduced in 2018 by Visentin et al. [40]. There are also computational models of glucose homeostasis that are liver-centric: linking the liver to other organ compartments [41, 42] with the goal of investigating how food composition influences hepatic lipid synthesis and could lead to the development of different types of diabetes [43, 44]. Despite their instrumental role, these compartment models often adopted a coarse-grained approach. In recent years, stoichiometry-based mechanistic models have become popular, employing a rate equation for each metabolic reaction within a cell or organ [29, 31–36]. For instance, Kurata [29] proposed a comprehensive whole-body model, incorporating enzyme and transporter reactions alongside hormonal regulation in postprandial and postabsorptive states. Panunzi et al. [45] extended a whole-body model initially proposed by Sorensen [46] by incorporating food intake. Sluka et al. [37] proposed a liver model for acetaminophen pharmacology, integrating three-scale modules of enzyme reactions within a cell, physiologically based pharmacokinetics of acetaminophen at organs, and its distribution at the whole-body level. Ashworth et al. [38] developed a spatial kinetic model of hepatic glucose and lipid metabolism, treating sinusoidal tissue units instead of single hepatocytes. Carstensen et al. [30] introduced a formalism for developing whole-body multi-scale models, based on constraint-based modeling approaches described by Yasemi and Jolicoeur [47]. Per their formalism, the metabolism inside the organs is explained by the stoichiometry of enzymatic reactions, and Michaelis-Menten kinetics are used to describe enzymatic reactions. Some other models have focused on the metabolism of energy expenditure, with Dash et al. [33] developing a computational model of skeletal muscle metabolism linking cellular adaptations induced by altered loading states to metabolic responses during exercise, and Kim et al. [31] developing a whole-body model of fuel homeostasis during exercise using hormonal control over cellular metabolic processes.

While many models provide valuable insights into metabolic processes, few adequately address the nuanced differences between male and female physiology. Palmisano et al. [23] and Abo et al. [20] developped sex-specific models of exercise with the aim of achieving greater generalization. Thiele et al. [14] introduced the virtual humans: Harvey and Harvetta, which are network stoichiometric reconstructions based on omics data that simulate steady-state metabolic fluxes. There is also *LiverSex*, a sex based multi-tissue and multi-level liver metabolic model [48]. Swapnasrita et al. [49] developed sex-specific models to compare kidney function in male and female patients with different stages of diabetes. This overview of the multi-compartment modelling literature is not an exhaustive tabulation of the diverse range of published metabolic models.

In this work, we introduce a sex-specific, multi-organ, and multi-scale whole-body model of metabolism, which accurately simulates key metabolite dynamics following various mixed meals. Six major organs—brain, heart, skeletal muscle, gastrointestinal (GI) tract, liver, and adipose tissue—are modeled, along with an *other tissues* compartment representing the rest of tissues. Metabolism within organs is modeled by stoichiometric enzymatic reactions. We used mass-action kinetics to describe these reactions. This model extends our previously published sex-specific model of energy metabolism during aerobic exercise [20]. Here, our goal is to connect cellular metabolism in organs with whole-body responses following different mixed meals, particularly high-carbohydrate and high-fat meals. We investigated sex-related differences in metabolic responses during both the absorptive and postabsorptive phases following a meal. We hypothesized that the female model will exhibit a greater reliance on lipid metabolism during both phases. Moreover, we hypothesized that whole-body sex differences will stem from organ-level variations, particularly in the liver, skeletal muscle, and adipose tissue, given the inherent differences in body composition between the sexes. Our objectives are to: (1) quantify sex differences in carbohydrate and lipid metabolism at the whole-body level, (2) assess sex differences in carbohydrate and lipid metabolism across various organs and tissues, identifying key organs driving whole-body responses, and (3) propose a candidate physiological mechanism driving sex differences in glucose production and fat oxidation patterns. Notably, point (3) addresses a gap in the experimental literature: while experiments demonstrate that increased hepatic free fatty acids (FFA) uptake and subsequent FFA oxidation enhance glucose production [50–52], women exhibit lower hepatic glucose output compared to men despite taking up and oxidizing more FFA [4, 11, 17, 53].

Novel contributions of our work to the existing literature involve developing a comprehensive, sex-specific model encompassing whole-body dynamics during feeding and fasting. Extending from our prior work on exercise metabolism, this computational model proves versatile, offering insights into various functions such as exercise, diet, and sex modulation.

The remainder of this paper is structured as follows: simulation results are presented in Section 2. In particular, Sections 2.1-2.2 provide an overview of model construction and results for model calibration and validation. Our main results are presented in Sections 2.3 and 2.4, covering whole-body and organ-specific metabolic responses, respectively. In Section 3, we discuss our key findings and relate them to model assumptions. Section 4 summarizes our results. Appendix A presents a detailed explanation of our mathematical model and modeling approach, while Appendix B outlines the parameter estimation process. A comprehensive list of all parameter values and model equations is available in S1 Supporting Information.

## 2 Results

### 2.1 Model construction

We present a whole-body model of food intake comprising seven compartments: brain, heart, liver, GI tract, skeletal muscle, adipose tissue, and “other tissues”, 22 metabolites and 25 reactions including the hormonal effect from pancreatic hormones, insulin and glucagon. *Other tissues* include kidneys, upper extremity muscles, and the rest of tissues. We specifically model the metabolic response to the intake of a mixed meal (carbohydrates and fat) from immediately after the meal (postprandial phase) to the postabsorptive phase (short-term fast). Figure 1 illustrates the multi-scale whole-body model. A fundamental assumption of the model concerning feeding and fasting states is that circulating levels of insulin and glucagon directly influence the carbohydrate and fat metabolism of organs and tissues, including the heart, skeletal muscle, GI tract, liver, and adipose tissue. Our model is tailored to sex, representing the metabolic states of healthy young adult man and woman. A complete mathematical description of the model is available in Appendix A.

**Fig 1.**
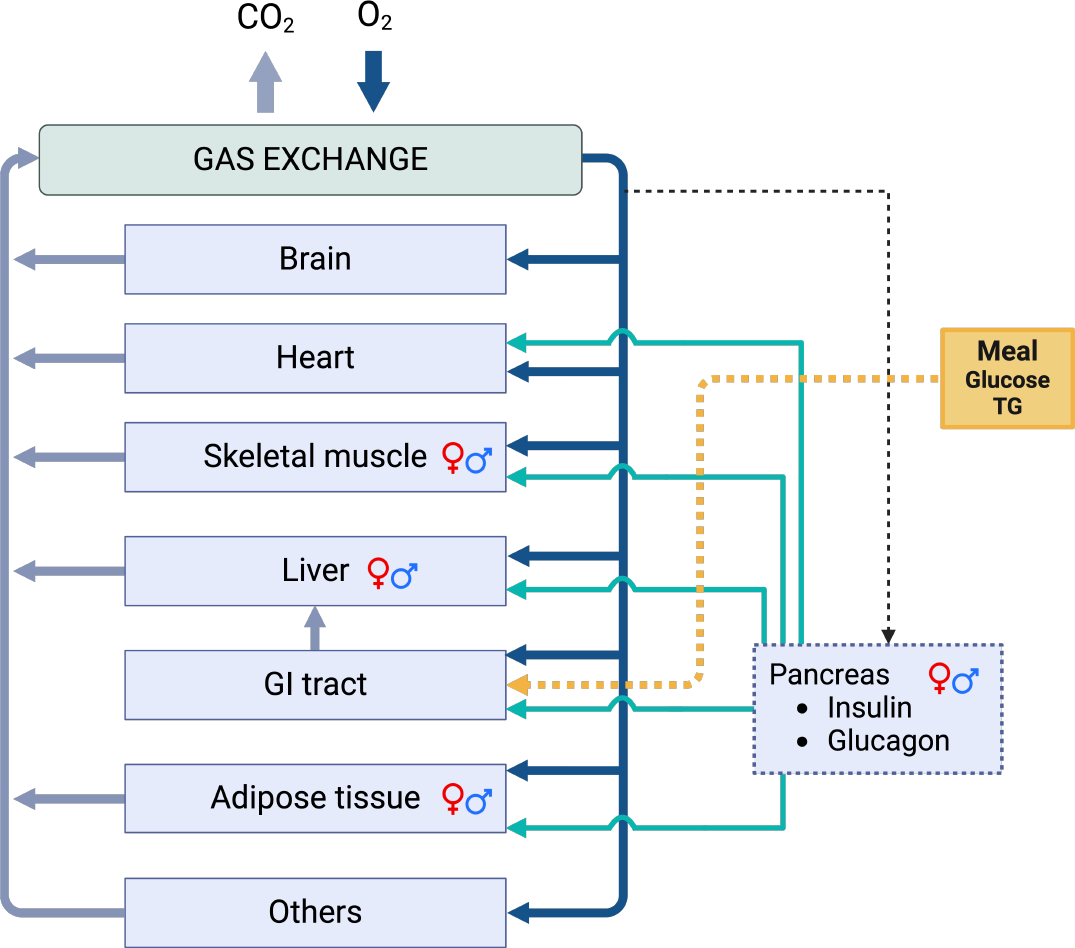
Whole-body system diagram. The systemic circulation connects all tissues/organs by transporting substrates in arterial oxygenated blood to the organs/tissues (solid dark blue arrows). Venous blood (solid light blue arrows) leaving these tissues/organs eliminates by-products and becomes arterial blood to restart circulation after releasing carbon dioxide and absorbing oxygen in the lungs (gas exchange). Blood supply to the liver comes from both the hepatic artery and venous blood from the GI tract. Nutrients are assimilated in the GI tract and subsequently enter the bloodstream. The nutrient-rich blood flows to all other organs and tissues. The pancreas responds to variations in arterial glucose concentration (indicated by the dashed arrow), regulating the levels of insulin and glucagon. Alterations in the concentrations of insulin and glucagon impact metabolic fluxes in the heart, skeletal muscle, liver, gastrointestinal tract, and adipose tissue (depicted by solid green arrows), thereby concluding the feedback regulatory mechanism. Male and female sex symbols represent compartments where sex-differences in reaction rates, besides differences in tissue/organ weights, are implemented.

### 2.2 Model calibration and validation

In this section, we show the time profiles of key metabolites following diverse single meals. Model simulations are presented alongside experimental data utilized for model calibration and validation. The results discussed here pertain to four experiments in which participants consumed a mixed meal after an overnight fast lasting 12 to 14 hours: (Experiment 1) 96 g carbohydrate and 33 g fat [54, 55], (Experiment 2) 139 g carbohydrate and 17 g fat [56], (Experiment 3) 58 g carbohydrate and 27.7 g fat [57], and (Experiment 4) 289 g carbohydrate and 45 g fat [58]. Specifically, selected sets of data from experiments 1, 2, and 4 were employed for model calibration, while data from experiments 2 (not used in calibration) and 3 were utilized to validate model predictions. We explored two metabolic states: (i) the absorptive or postprandial state, also known as the fed state, is defined as the period (0–6h) following meal ingestion, encompassing the processes of nutrient digestion and absorption; (ii) the postabsorptive state (6–12h), a fasting period that involves the utilization and storage of nutrients in specific tissues [59].

To formulate the male model, we first identified parameter values (see Table 1) using data from experiments 1, 2, and 4 [54–56, 58]. To formulate the female model, we extended our previously published model that simulates a woman’s metabolism during aerobic exercise to simulate a meal [20]. The meal-related model components and parameters were taken to be the same as in the male model. We then validated both models by comparing their predictions with additional data from experiments 2 and [56, 57]. We emphasize that all simulations for the female model are predictions.

**Table 1.** General description of estimated model parameters.

|  | Parameters | Description | Equations |
| --- | --- | --- | --- |
| Insulin | $k_I^0$ | basal secretion rate | (7) |
| | $V_{\max_I}$ | maximum rate coefficient | (7) |
| Glucagon | $k_G^0$ | basal secretion rate | (8) |
| | $V_{\max_G}$ | maximum rate coefficient | (8) |
| | $n_2, n_3$ | hill coefficients | (8) |
| Dietary input | $K_G$ | half-emptying glucose threshold | (5) |
| | $K_T$ | half-emptying fat threshold | (6) |
| (Post)absorptive regulation | $V_{\max_\beta}^x, V_{\max_\gamma}^x$<br>$V_{\max_{\beta,\text{fat}}}^x, V_{\max_{\gamma,\text{fat}}}^x$ | maximum rate coefficients for enzyme activity factors $(B_x, \Gamma_x, B_x^{\text{fat}}, \Gamma_x^{\text{fat}})$ | (18-25) |
| | $K_{\max_\beta}^x, K_{\max_\gamma}^x$<br>$K_{\max_{\beta,\text{fat}}}^x, K_{\max_{\gamma,\text{fat}}}^x$ | Michaelis constants for enzyme activity factors $(B_x, \Gamma_x, B_x^{\text{fat}}, \Gamma_x^{\text{fat}})$ | (18-25) |
| | $n, k, m, q$ | hill coefficients for enzyme activity factors $(B_x, \Gamma_x, B_x^{\text{fat}}, \Gamma_x^{\text{fat}})$ | (18-25) |

#### 2.2.1 Postprandial dynamics of insulin and glucose

To calibrate the model, model parameters were chosen such that the model adequately simulates the temporal evolution of key metabolites after an overnight fast and following a single meal. Figure 2 shows the time course of plasma insulin and plasma glucose for the male model. Similar profiles are observed for the female model (data not shown). Following a meal, glucose enters the bloodstream by absorption from the intestine. In general, a rise in blood glucose concentration becomes apparent within approximately 15 minutes and reaches its peak around 30–60 minutes after a meal, subsequently returning to the baseline level of approximately 5 mM roughly two hours post-meal. Similar timescales of exogenous glucose appearance into the bloodstream have been reported in the literature [29, 54, 55, 59]. Our results indicate no sexual dimorphism in the glucose and insulin responses to a mixed meal (data not shown) during the absorptive (0–6h) and early postabsorptive phases (6–12h). The absence of sexual dimorphism was also reported in a study involving young and healthy individuals during the absorptive phase [15]. Regarding the postabsorptive phase, some studies have shown no significant sex difference between women and men for fasts of up to 22 hours, where individuals were either matched on percent body fat [17] or not [11].

**Fig 2.**
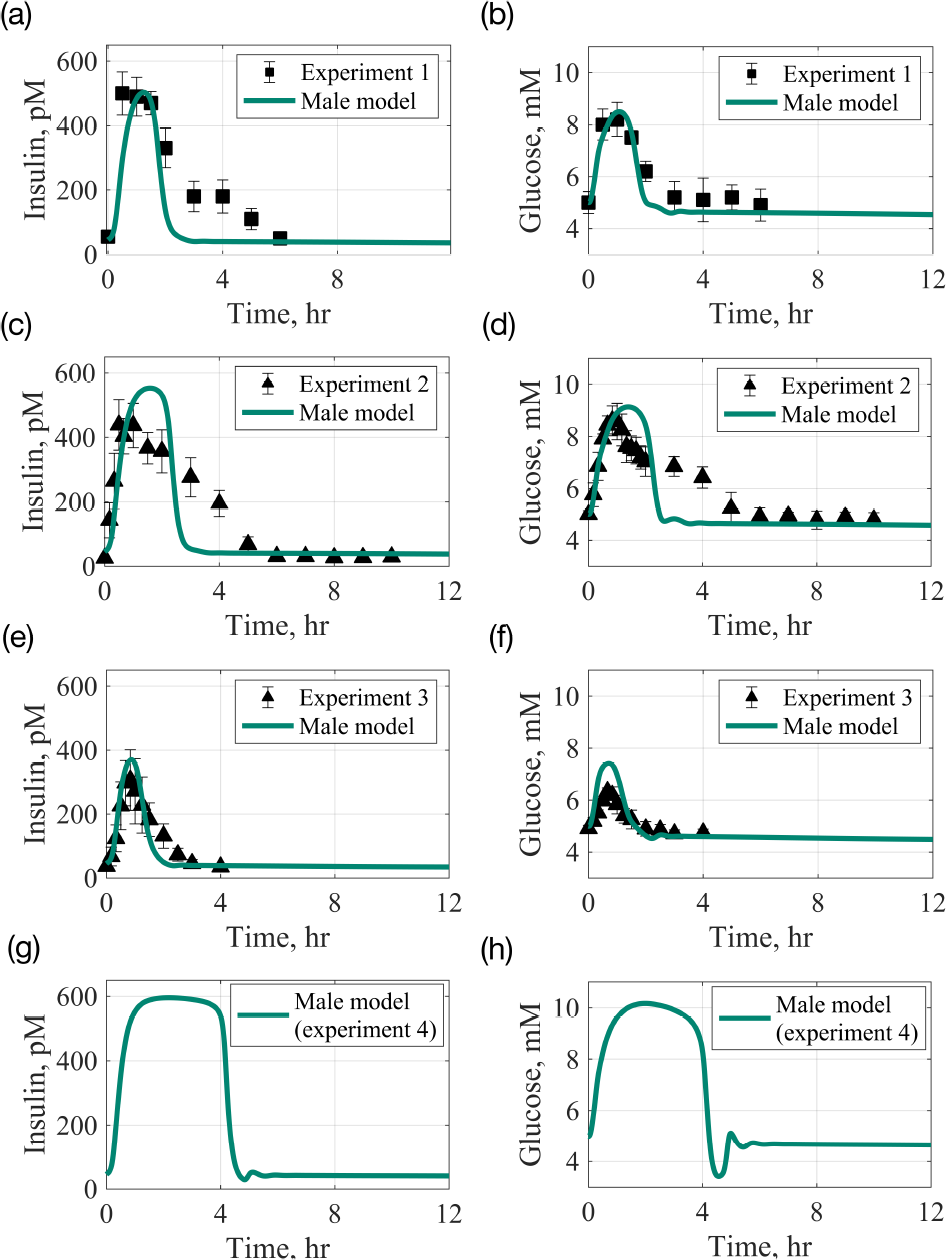
Time profile of plasma insulin and glucose concentrations after an overnight fast and following a single meal. Experiment 1: 96 g carbohydrate and 33 g fat [54]; Experiment 2: 139 g carbohydrate and 17 g fat [56]; Experiment 3: 58 g carbohydrate and 27.7 g fat [57]; Experiment 4: 289 g carbohydrate and 45 g fat [58]. Square markers (▪) with lines represent calibration data with standard errors [54]; Triangular markers (▴) with lines represent validation data with standard errors [56, 57]. Lines represent model simulations. (a), (c), (e), (g): plasma insulin; (b), (d), (f), (h): plasma glucose. Absorptive phase, 0–6h; postabsorptive phase, 6–12h.

Using the parameters identified above, we conducted a comparison between model predictions and data from experiments 2 and 3 that were not used in the model calibration [56, 57]. In Figs 2c-2f, the time profiles of insulin and glucose are presented after a single meal containing 139 g of carbohydrates and 17 g of fat (experiment 2, data from Ref. [56]), as well as after a single meal with 58 g of carbohydrates and 27.7 g of fat (experiment 3, data from Ref. [57]). Our simulations exhibit good qualitative agreement with the experimental data; however, certain discrepancies are noted. Specifically, our model tends to overestimate the insulin peak in Fig 2c and the glucose peak in Fig 2f. For the former (Fig 2c), the rise in glucose is similar to that observed in our calibration dataset, resulting in insulin reaching a level comparable to what was observed in the calibration dataset (Fig 2a). The type and composition of a meal influence insulin secretion. Meals with higher carbohydrate content (Fig 2c-d, 139g of carbohydrates) are expected to yield higher insulin excursions compared to meals with lower carbohydrate content (Fig 2a-b, 96g of carbohydrates). However, it is important to note that the glycemic index of a meal, contributing to different insulin responses, could introduce variability into the observations [60]. In the case of the glucose peak in Fig 2f, as the dataset does not explicitly specify the proportion of carbohydrate represented by glucose, we assumed that the entire 58g corresponds to glucose. This assumption may account for the discrepancy observed in our simulations, where the glucose peak is approximately 7.5 mM compared to the dataset’s peak of around 6.5 mM. It is worth noting that the nature and composition of a meal can influence the elevation of glucose levels [60].

Post-meal hyperglycemia and glucose clearance depend on three factors: gut-emptying times, peak times, and the time-to-return to baseline concentrations for glucose and insulin. We provide a detailed explanation of how we model gut-emptying times in Appendix A, Eqs 3-6. These factors are largely influenced by the size and composition of the meal. Fig 2 illustrates that meals with carbohydrate quantities of 58g (experiment 3), 96g (experiment 1), 139g (experiment 2), and 289g (experiment 4) have gut-emptying times for glucose of 39, 50, 61, and 79 minutes, respectively. Trends for peak times and times-to-return to baseline are similar: larger carbohydrate loads lead to delayed glucose peaks, with substantial meals causing postprandial hyperglycemia for over 3 hours. Notably, in the case of experiment 4 (289 g of carbohydrates), glucose peaks after 1 hour and remains elevated for over 3 hours after the meal. Such an occurrence characterizes postprandial hyperglycemia, formally defined as plasma glucose levels exceeding 7.8 mmol/L 1-2 hours after food intake, and can lead to vascular complications, particularly in diabetic patients [61]. For larger meals, like those in experiments 2 and 4 shown in Fig 2, an undershoot occurs as glucose concentration returns to baseline. This happens because insulin usually peaks and returns to its baseline levels 10-30 minutes after glucose. Therefore, a large carbohydrate meal prolongs elevated circulating insulin levels, causing a delay in the time organs stay sensitive to insulin and, in turn, uptake glucose. This delay manifests as an initial undershoot in glucose levels before eventually stabilizing at basal levels.

Overall, we observe no sexual dimorphism in insulin and glucose responses following different mixed meals, and the simulated results are consistent with experimental data from several healthy subjects [54, 56, 57, 59].

#### 2.2.2 Postprandial dynamics of glycogen storage

In the postprandial phase, glucose is stored as glycogen through a process called glycogenesis, primarily occurring in the liver and skeletal muscle [59]. Figure 3 shows the change in glycogen, relative to its initial concentration, in liver (left column) and skeletal muscle (right column). The female model tends to accumulate more glycogen than the male model in both organs, particularly after a carbohydrate-rich meal (*>*100 g carbohydrate, liver: Figs 3c and 3g; skeletal muscle: Figs 3d and 3h). We note that basal glycogen concentrations are the same for both organs in both models, as indicated in the literature [62, 63]. In these organs, glycogen concentrations increase due to insulin action following a meal (i.e., insulin activates glycogenesis), as confirmed by experimental data [56, 58]. The relative change in hepatic glycogen stores and the duration until the initiation of glycogen breakdown for energy needs depend on the size of the carbohydrate load. Specifically, for meals comprising 58g, 96g, 139g, and 289g of carbohydrates, glycogen concentrations in the male model increase by 15%, 21%, 33%, and 80%, respectively, peaking at 1.5, 2, 3, and 4.5 hours, respectively. In the female model, glycogen concentrations rise by 15%, 27%, 42%, and 100%, respectively, with similar peak times as the male model. Although the peak times in skeletal muscle align with those in the liver, the extent of glycogen accumulation is lower in skeletal muscle for both sexes. We observed increases of 3%, 5%, 7%, and 14% in the male model and increases of 5%, 8%, 12%, and 21% in the female model following carbohydrate loads of 58g, 96g, 139g, and 289g, respectively. These results imply a preferential replenishment of hepatic glycogen stores over intramuscular stores.

**Fig 3.**
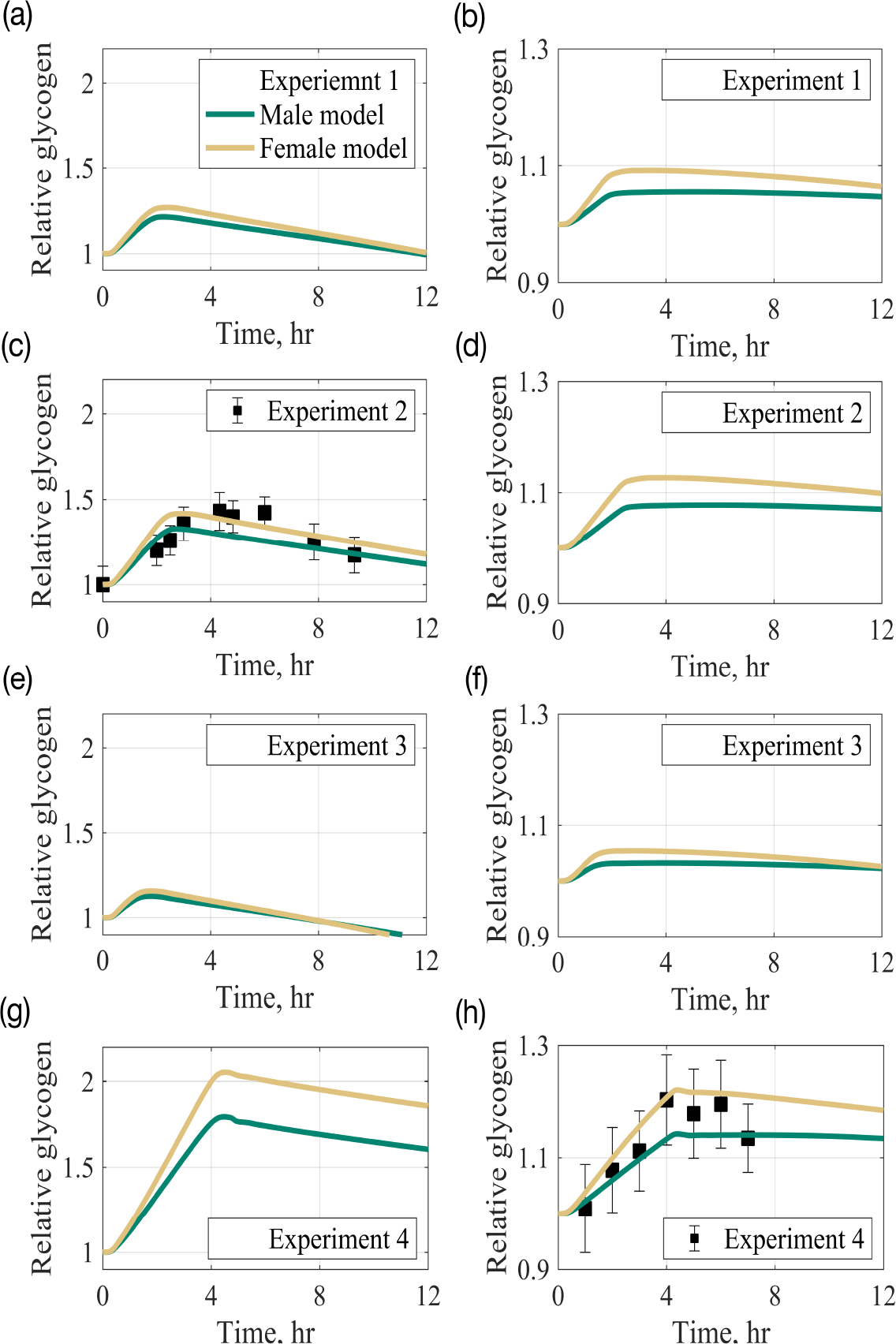
Time profile of glycogen concentration in liver (left column) and skeletal muscle (right column), relative to its initial value, after an overnight fast and following a single meal. Experiment 1: 96 g carbohydrate and 33 g fat [54]; Experiment 2: 139 g carbohydrate and 17 g fat [56]; Experiment 3: 58 g carbohydrate and 27.7 g fat [57]; Experiment 4: 289 g carbohydrate and 45 g fat [58]. Square markers (▪) with lines represent calibration data with standard errors [56, 58]. Lines represent model simulations.

A few hours after the meal, liver glycogen undergoes degradation into glucose (glycogenolysis), which is then released into the bloodstream. Our model simulations demonstrate relatively constant blood glucose levels during the postabsorptive phase (beyond 6 hours; see Fig 2, right column), attributed to liver glycogen being degraded into glucose via glycogenolysis and subsequently released into the bloodstream.

Skeletal muscle glycogen remains relatively constant during the postabsorptive phase, as indicated by Ref. [64]. This observation holds true irrespective of the meal composition and sex, as shown in the right column of Fig 3. Skeletal muscle lacks the essential enzyme glucose 6-phosphatase for glycogenolysis and therefore cannot release glucose. Muscle glycogen primarily serves as a local energy substrate for exercise rather than as an energy source to maintain blood glucose concentration during fasting [64]. The body’s major fuel store is TG in adipose tissue, while liver and muscle glycogen serve as short-term carbohydrate stores [59].

#### 2.2.3 Postprandial dynamics of other key metabolites

Figure 4 shows the time evolution of plasma lactate, TG, FFA, and glycerol concentrations after an overnight fast and following a single meal of 96g carbohydrate and 33g of fat (Experiment 1). Model predictions show consistent trends across various meal compositions for both female and male models (data not shown). Beside glucose, there is also an elevation of the plasma lactate concentration after ingestion of carbohydrates (Fig 4a), as indicated in [54]. There is qualitative agreement between simulations and experimental data for plasma lactate, although the peak in the simulations occurs approximately two hours after the experimentally observed peak. While the increase in blood glucose and lactate production can be correlated, they may not peak at the exact same time. Lactate levels are influenced by various factors, including tissue-specific metabolic activities, oxygen availability, and the overall metabolic state of the body [65–67]. For instance, Ref. [66] showed that following the uptake of carbohydrates, plasma lactate, glucose, and insulin increased within 15-30 min, reaching peak levels at 180, 90, and 90 min, respectively. Numerical simulations show no sex-related differences in lactate response following a mixed meal.

**Fig 4.**
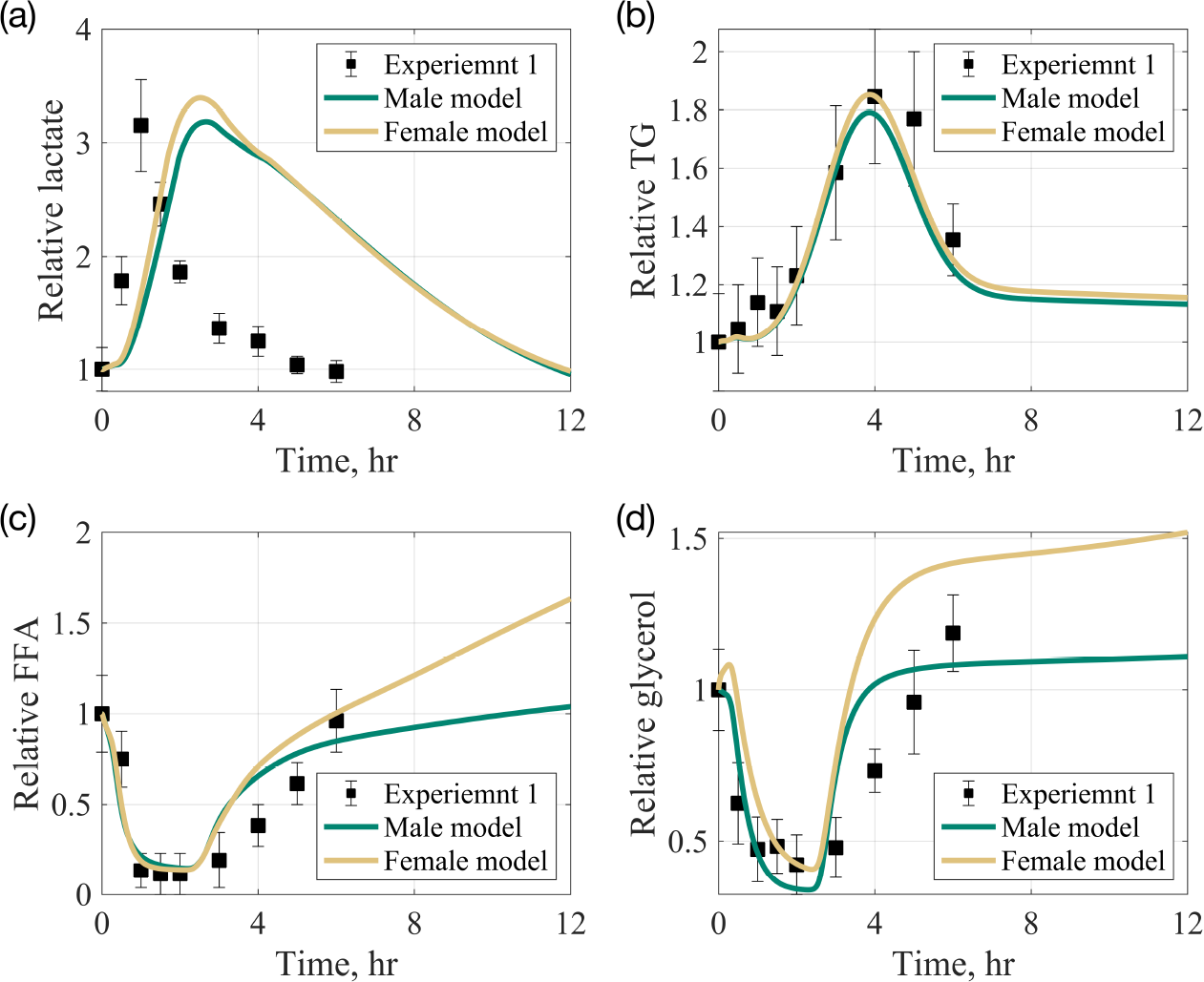
Time profile of plasma metabolite concentrations after an overnight fast and following a single meal. Experiment 1: 96 g carbohydrate and 33 g fat. Square markers with lines represent experimental data with standard errors [54, 55]; Lines correspond to the male and female model simulations. (a)-(d) concentrations, relative to initial values, of plasma lactate, plasma free fatty acids (FFA), plasma triglycerides (TG), and plasma glycerol, respectively. Basal concentrations of the following substrates differ significantly between the sexes: glucose (5 vs. 4.91 mM) [16], FFA (0.66 vs. 0.76 mM) [16], and TG (0.99 vs. 0.93 mM) [68] in males and females, respectively. The initial concentrations of other substrates are taken to be the same in both male and female models. Absorptive phase, 0–6h; postabsorptive phase, 6–12h.

Dietary fat is absorbed at a much slower rate than glucose, so the peak in plasma TG concentration occurs 4 hours after the meal (Fig 4b). Following the initial post-meal peak, plasma TG gradually decreases, in line with experimental data [5]. Directly after the meal, plasma FFA and glycerol drop, then gradually increase to pre-meal levels after approximately 4 hours (Fig 4c and Fig 4d). This decrease may be caused by the reduced release of FFA (and glycerol) from adipose tissue into the bloodstream. The increase in insulin directly suppresses lipolysis in adipose tissue, causing a postprandial reduction in plasma FFA and glycerol concentrations, as evidenced in the literature [59]. General lipid responses (TG, FFA, glycerol) to a mixed meal are sex-neutral in the postprandial phase. However, sexual dimorphism becomes evident as soon as 4 hours after the meal, particularly for FFA and TG. The surge in FFA and glycerol, the products of lipolysis, is greater in the female model than in the male model. Similar sex differences in lipid metabolism, but not glucose metabolism, have also been reported in the literature [17, 53].

### 2.3 Whole-body sexual dimorphism emerge in lipid metabolism

The respiratory quotient (RQ) reflects the relative oxidation levels of macronutrients to meet metabolic demands. Precisely, RQ is the metabolic exchange of gas ratio at the cellular level that equals the ratio of CO_2_ eliminated to oxygen consumed. *In vivo*, it is directly measured from blood, while *in silico*, we use the ratio between CO_2_ release rate into the blood compartment and oxygen uptake rate across all tissues/organs [20]. This method provides an indirect calorimetry measure indicating the primary fuel source (e.g., carbohydrate or fat) supporting the body’s energy needs [69]. Physiological RQ values typically range between 0.7 and 1.0, varying with substrate oxidation; glucose has a RQ of 1.0, while fat has a RQ of 0.7 [70]. RQ inversely correlates with lipid oxidation, where a high RQ signifies low lipid oxidation and high carbohydrate oxidation.

Figure 5 illustrates the dynamics of whole-body RQ during both the absorptive (0-6h) and postabsorptive (*>*6h) phases for two meal compositions: high-carbohydrate and high-fat meals (refer to Table 2). Irrespective of sex and meal composition, there is a significant increase in RQ during the absorptive phase (RQ*≥*0.88), indicating a predominant utilization of carbohydrates for energy metabolism. In the absorptive phase, female RQ shows a higher increase than male RQ, suggesting a slightly higher reliance on carbohydrate oxidation in the female model. As the absorptive phase transitions to the postabsorptive phase, a general decrease in RQ is observed. In the postabsorptive phase, RQ values remain elevated (RQ*>* 0.8), but no discernible differences in RQ between sexes become apparent. These RQ values indicate sustained carbohydrate oxidation during the early postabsorptive phase.

**Table 2.** Meal compositions.

| Meal type | % CHO | % Fat | CHO (g) | Fat (g) | CHO (cal) | Fat (cal) |
| --- | --- | --- | --- | --- | --- | --- |
| High-carbohydrate (HiC) | 90 | 10 | 180 | 8.9 | 720 | 80 |
| High-fat (HiF) | 50 | 50 | 100 | 44.4 | 400 | 400 |

We consider isocaloric meals consisting of 800 calories (kcal). Fat (lipids) provides 9 kcal/g, whereas carbohydrates offer approximately 4 kcal/g [59].

**Fig 5.**
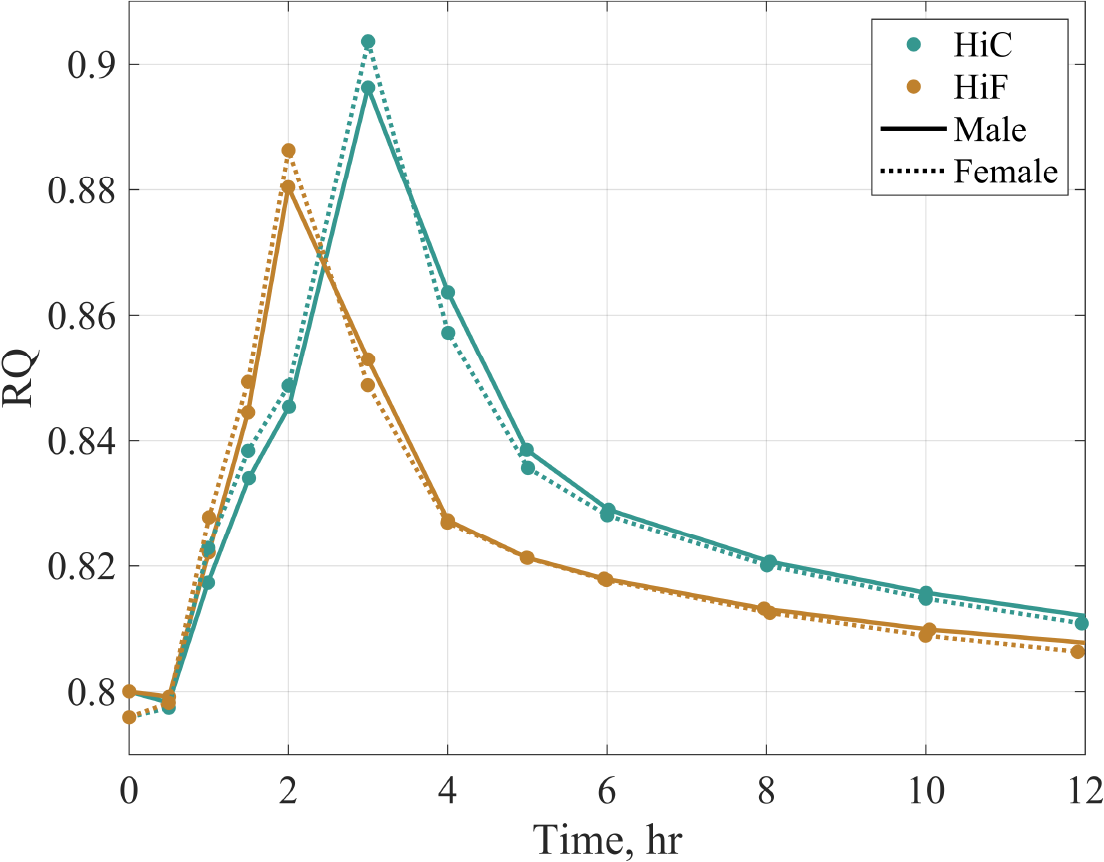
Time profile of whole-body respiratory quotient (RQ) in response to a single 800 kcal meal. Two distinct meal types were investigated: high-carbohydrate (HiC) and high-fat (HiF) meals. The whole-body RQ was calculated as the ratio of 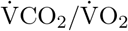, where 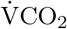and 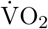 represent the sums of CO_2_ production and O_2_ consumption rates across all organs and tissues, respectively. We assumed that the respiratory exchange ratio (RER) reflects systemic nonprotein RQ, as suggested in Ref. [71]. Absorptive phase, 0–6h; postabsorptive phase, 6–12h.

In both male and female models, RQ reaches its peak 2 hours after a high-fat meal and 3 hours after a high-carbohydrate meal. This observation, where RQ peaks earlier for a high-fat meal compared to a high-carbohydrate meal, is not trivial and may be linked to a differential modulation of metabolic pathways by these macronutrients. Figure 6 shows the fractions of carbohydrate and fat utilized for energy production, specifically for ATP hydrolysis (*ϕ*_ATP*→*ADP_). There is no noticeable difference between the sexes in the oxidation fractions. However, once again, the peak in carbohydrate oxidation occurs earlier for the high-fat meal compared to the high-carbohydrate meal (Fig 6a). For the high-fat meal, carbohydrate oxidation peaks after 2 hours and remains elevated until hour 4. In contrast, for the high-carbohydrate meal, carbohydrate oxidation peaks after 3 hours and remains elevated until hour 6. Fat oxidation, significantly reduced during the absorptive phase, also exhibits a similar difference in trough time between the different meal compositions (Fig 6b). During the postabsorptive phase, the contribution of fat to energy production increases to exceed the contribution of carbohydrates. This transition from carbohydrate to fat metabolism results in an eventual decline in RQ, as shown in Fig 5.

**Fig 6.**
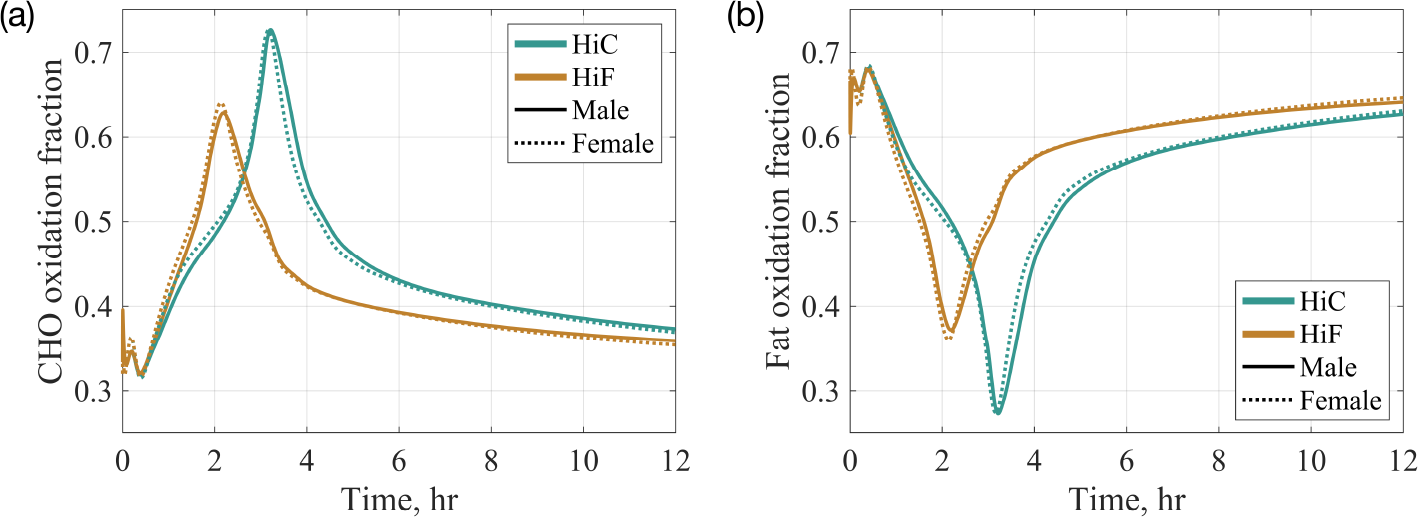
Carbohydrates and fat oxidation fractions in response to a single 800 kcal meal. HiC, high-carbohydrate meal; HiF, high-fat meal; CHO, carbohydrate. The oxidation fractions (unitless) establish a relationship between RQ values and the actual proportion of carbohydrates and fat utilized for ATP hydrolysis (*ϕ*_ATP*→*ADP_). Indirect calorimetry methods, as outlined by Roepstorff et al. [71], were employed: CHO oxidation fraction = (RQ - 0.7)/0.3; fat oxidation fraction = 1 - CHO oxidation fraction. Absorptive phase, 0–6h; postabsorptive phase, 6–12h.

A closer examination of whole-body metabolic fluxes (Table 3) reveals sexual dimorphism during both the absorptive and postabsorptive phases. We introduced a quantity called the percent relative difference between the sexes (Δ_*F/M*_). For each flux, it is calculated as (female flux/male flux *−* 1) *×* 100. During the absorptive phase and for both diet types, the female model exhibits greater rates of glycogen storage (*>* 5%) compared to the male model, along with similarly higher rates of *de novo* lipogenesis (*>* 5%)—the conversion of excess carbohydrate into fatty acids that are then esterified and stored as TG. Given that blood glucose levels are similar between the sexes during the absorptive phase (Fig 2), this result suggests that the higher rate of net glycogenolysis may not directly contribute to raising blood glucose. Instead, a higher amount of carbohydrates may be redirected for storage as fat in the female model rather than being utilized for direct oxidation. Moreover, the female model exhibits lower rates of TG breakdown (*< −*20%) compared to the male model. Overall, the sex-related difference is more prominent in fat metabolism than glucose metabolism immediately after a meal. Thus, metabolic changes at the whole-body level in the absorptive phase could involve sexual dimorphism in lipid but not glucose metabolism, as indicated in Ref. [15].

**Table 3.** Whole-body metabolic fluxes.

| Metabolic flux (mmol/min)<sup>a</sup> | HiC |  |  | HiF |  |  |
| --- | --- | --- | --- | --- | --- | --- |
| | Male | Female | $\Delta_{F/M}$ | Male | Female | $\Delta_{F/M}$ |
| Absorptive phase |  |  |  |  |  |  |
| Net glycogen breakdown<sup>b</sup> | -1.05 | -1.11 | 5.25% | -0.708 | -0.753 | 6.32% |
| Net gluconeogenesis<sup>c</sup> | -1.01 | -1.05 | 4.06% | -0.791 | -0.814 | 2.90% |
| Net TG breakdown<sup>d</sup> | 0.041 | 0.0321 | **-21.7%** | 0.0427 | 0.0314 | **-26.5%** |
| *De novo* lipogenesis<sup>e</sup> | 0.0193 | 0.0204 | 5.47% | 0.0177 | 0.0186 | 5.37% |
| Postabsorptive phase |  |  |  |  |  |  |
| Net glycogen breakdown | 0.335 | 0.372 | **10.8%** | 0.347 | 0.366 | 5.42% |
| Net gluconeogenesis | -0.380 | -0.320 | **-15.8%** | -0.309 | -0.311 | 0.902% |
| Net TG breakdown | 0.0936 | 0.101 | **8.31%** | 0.0954 | 0.103 | **7.64%** |
| *De novo* lipogenesis | 0.0125 | 0.0127 | 1.71% | 0.0127 | 0.0132 | 3.40% |

<sup>a</sup> Values are averages over the specified phase and following a single 800 kcal meal. HiC, high-carbohydrate meal; HiF, high-fat meal. Percent change in the female model relative to the male model ( $\Delta_{F/M}$ ) for each flux is calculated as (female flux/male flux $-1$ ) × 100.
<sup>b</sup> Net glycogen breakdown is defined as the difference in metabolic rates between glycogenolysis (breakdown of glycogen) and glycogenesis (production of glycogen).
<sup>c</sup> Net gluconeogenesis is defined as the difference between gluconeogenesis II (production of glucose-6-phosphate), specifically in the liver, and glycolysis II (utilization of glucose-6-phosphate).
<sup>d</sup> Net TG breakdown is defined as the difference between lipolysis (TG breakdown) and TG synthesis.
<sup>e</sup> *De novo* lipogenesis refers to the conversion of glyceraldehyde-3-phosphate to glycerol-3-phosphate.
All fluxes, except for *de novo* lipogenesis, are net fluxes for which a positive value indicates breakdown (or release) of the associated substrate, and a negative value indicates storage (or utilization). Absorptive phase, 0–6h; postabsorptive phase, 6–12h.

Interestingly, the trend in the female model for fat preservation is reversed during the postabsorptive phase for both high-fat and high-carbohydrate meals, where the female model shows an approximately 8% higher net rate of TG breakdown compared to the male model. Overall, sexual dimorphism is more prominent in the postabsorptive phase following a high-carbohydrate meal. For instance, the net rate of glycogen breakdown is 10% higher, and the net rate of gluconeogenesis is 15% lower in the female model. Since glucose output in the postabsorptive phase is a combination of glycogen breakdown and gluconeogenesis, with these processes contributing almost equally to glucose production (see net rates in Table 3; postabsorptive), there is a net decrease of about 5% in the rate of glucose output in the female model compared to the male model. Consequently, the observed increase in systemic fat oxidation (*>*8%) for the female model may compensate, in part, for the decrease in glucose availability.

It is worth noting that the net rate of systemic gluconeogenesis is negative in both the absorptive and postabsorptive phases. This is intuitive during the absorptive phase: all organs and tissues are utilizing glucose (*glycolysis*) at a faster rate than the liver performs gluconeogenesis (marginal during this phase). However, during the postabsorptive phase, this suggests that the rate of glucose synthesis (hepatic gluconeogenesis) is lower than the rate of glycolysis by other organs. As a result, blood glucose levels decrease during the postabsorptive phase (refer to Fig 4). This aligns with our model assumption that blood glucose decreases at rest at a rate of 0.03 mmol/min.

### 2.4 Sexual dimorphism is tissue-specific and differences start appearing even during short-term fasting

#### 2.4.1 Tissue-specific sex differences in the absorptive phase

In the absorptive phase, ingested carbohydrates elevate blood glucose levels. Insulin is released, promoting glucose uptake by cells for energy or storage as glycogen. Simultaneously, dietary fats are broken down into FFA, absorbed into the bloodstream, re-esterified, and stored as TG within cells. These processes ensure energy balance and nutrient storage after a meal, contributing to maintaining blood glucose homeostasis and regulating lipid storage in the body. In this section, our focus is on exploring potential sex-related variations in substrate storage and utilization, as well as the potential implications of these differences for metabolic cross-talk among organs and tissues.

Glycogen, the storage form of glucose, is primarily stored in the liver and skeletal muscle. While other organs, such as the heart and small intestine, contain glycogen in smaller amounts, the liver and skeletal muscle are the main reservoirs. Figure 7 shows the change in glycogen concentration in the liver (Fig 7a) and skeletal muscle (Fig 7b) after a single meal, comparing high-carbohydrate and high-fat content meals.

**Fig 7.**
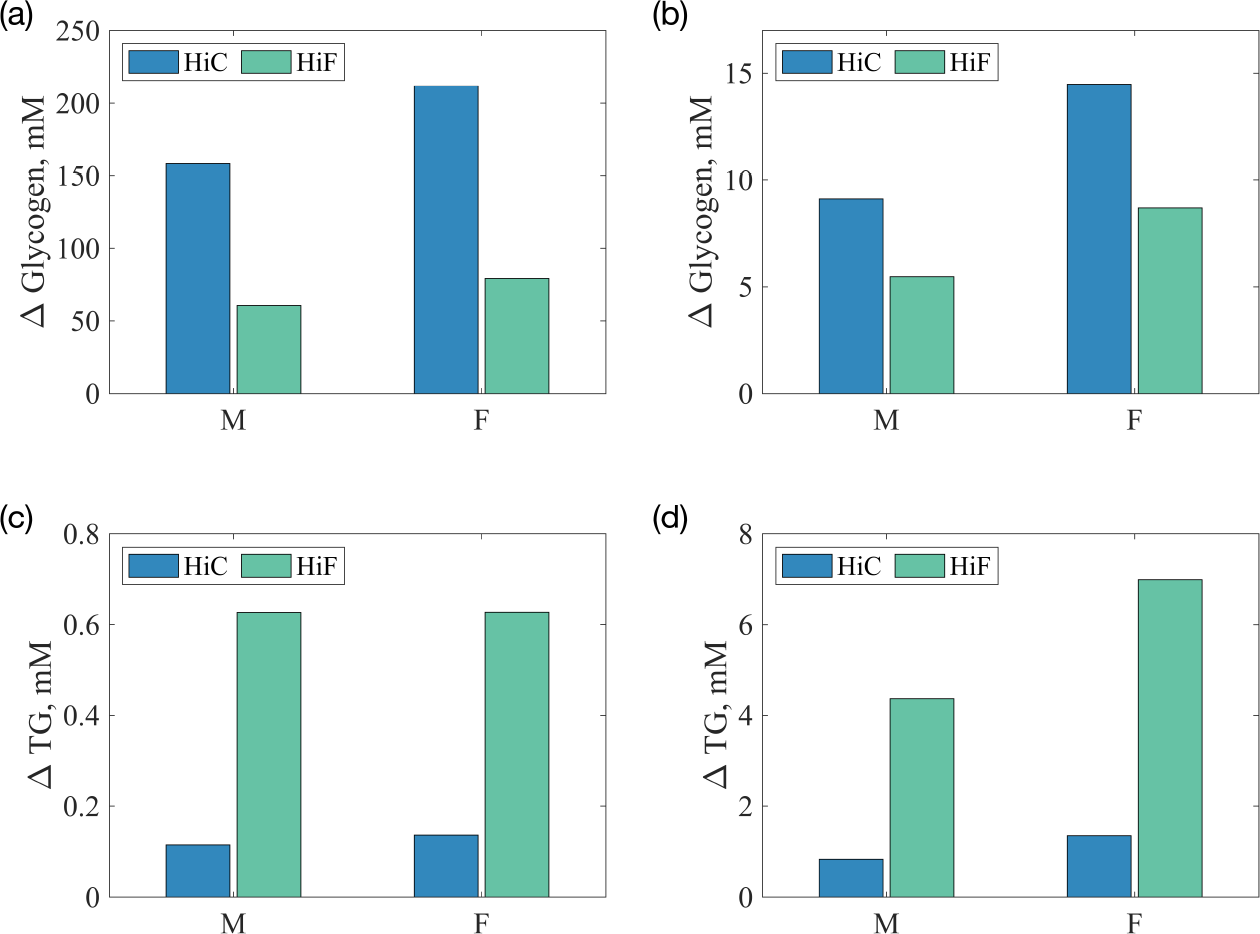
Carbohydrates (glycogen) and fat (TG) storage during the absorptive phase (0–6h). A single meal of 800 kcal is simulated at t=0. (a) liver glycogen; (b) skeletal muscle glycogen; (c) liver TG; (d) skeletal muscle TG. HiC, high-carbohydrate meal; HiF, high-fat meal. Δ refers to the absolute change in a given substrate, *C*_*x,i*_(*T*) *− C*_*x,i*_(0), where *C*_*x,i*_ is the concentration of substrate *i* in tissue *x*, and *T* = 6h.

Irrespective of sex, the liver accumulates more glycogen compared to skeletal muscle during the absorptive phase. In particular, the liver stores about 15 times more glycogen following a high-carbohydrate meal and about 9 times more glycogen following a high-fat meal than skeletal muscle. A comparison between sexes shows that the female model stores more glycogen in both the liver and skeletal muscle compared to the male model, a trend consistent across diet types. Specifically, the female model stores approximately 30% more hepatic glycogen and 58% more intramuscular glycogen than the male model, following either meals. Unlike glycogen, which is a short-term energy storage molecule in the liver and muscles, TG are primarily stored in adipose tissue, providing a concentrated and long-lasting reservoir of energy. However, TG are also stored in small amounts in other tissues, including the liver and skeletal muscle. Regarding hepatic TG, our model shows marginal differences in TG concentration between the sexes during the absorptive phase (Fig 7c). However, the female model stores approximately 60% more intramuscular TG, termed intramyocellular lipids (IMCL), than the male model following either meals (Fig 7d). Experimental studies have shown that women store more IMCL than men [3, 8]. Women were found to have 84% higher lipid density than men [8]. Our predictions agree well with quantitative experimental findings, as shown in Fig 7d.

#### 2.4.2 Tissue-specific sex differences in the postabsorptive phase

In our models, sex-specific differences become evident during short-term fasting, also referred to as short-term calorie restriction. We thus investigated sex-related differences in carbohydrate and fat metabolism during a short-term fast of 24 hours following a single 800 kcal meal (see Fig 8).

**Fig 8.**
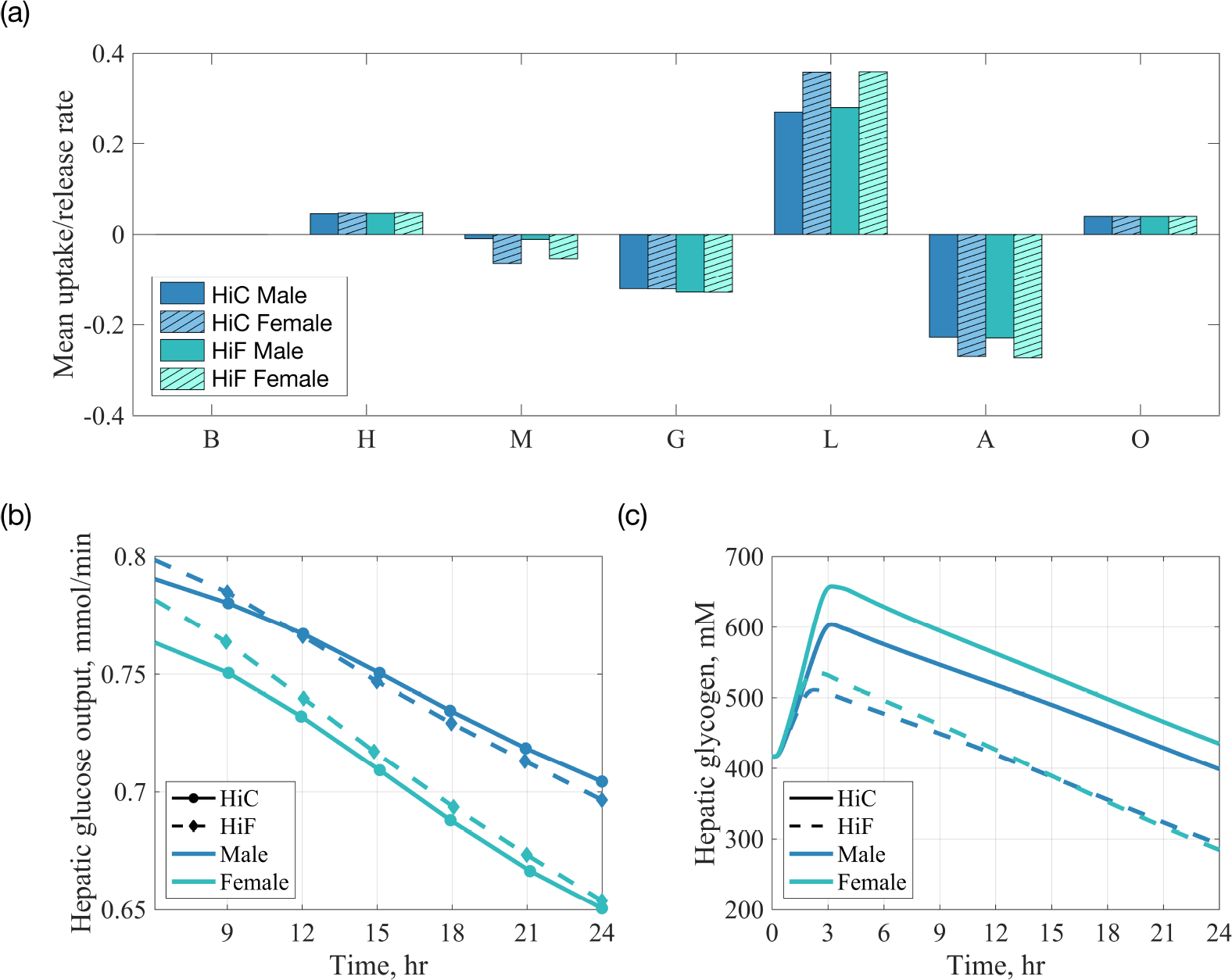
Metabolism of carbohydrates during a short-term fast (24h) following a single 800 kcal meal. HiC, high-carbohydrate meal; HiF, high-fat meal. (a) average uptake (or release) rate (mmol/min) of FFA per organ during the last 12 hours of the fasting window following HiC and HiF meals; (b) rate of hepatic glucose output (mmol/min) into the blood; (c) concentration of hepatic glycogen; mM, mmol/L. B, brain; H, heart; M, muscle; G, GI tract; L, liver, A, adipose tissue; O, other tissues.

Figure 8a shows the uptake and release rates of FFA by organs and tissues during the last 12 hours of a 24-hour fast. FFA flux is nil for the brain compartment as the blood-brain barrier restricts the passage of large, hydrophobic molecules like FFA. Instead, the brain predominantly relies on other energy sources, primarily glucose. For both sexes, adipose tissue and the GI tract release the most FFA, while the liver uptakes the most FFA. Although no significant sex difference appears for the GI tract, skeletal muscle, liver, and adipose tissue exhibit sex-related differences. For both high-carbohydrate and high-fat meals, the release (and uptake) rates of FFA are higher in the female model compared to the male model. In particular, female skeletal muscle releases 6 times more FFA during the fast period following the high-carbohydrate meal and 5 times more FFA following the high-fat meal. Female adipose tissue releases 17% and 21% more FFA during the fast period following the high-carbohydrate and high-fat meals, respectively. Consequently, the female liver uptakes FFA at rates 32% and 28% higher than the male model for high-carbohydrate and high-fat meals, respectively. Overall, more FFA is being released into the circulation during the fast than is being taken up by tissues in the female model, thus explaining the higher increase in plasma FFA in the postabsorptive phase in the female model compared to the male model (refer to Fig 4c), as observed in experiments [11, 17].

The increased reliance of the female liver on circulating FFA may be attributed to intracellular sex differences in substrate handling. For instance, as illustrated in Fig 8b, hepatic glucose production is lower in the female model compared to the male model during a short-term fast. Moreover, the difference in glucose release between the sexes becomes larger over time. This result applies to both high-carbohydrate and high-fat meals. Furthermore, Fig 8c shows that, alongside storing more glycogen during the absorptive phase, the female liver tends to maintain higher concentrations of glycogen during short-term fasting compared to the male model, specifically following a high-carbohydrate meal. There is no sex-related difference in hepatic glycogen content after a high-fat meal, yet the difference in hepatic glucose production persists. The tendency of the female liver to retain more glycogen throughout both absorptive and postabsorptive phases could explain the reduced glucose output, and thus the stronger reliance on FFA, potentially as a buffer to meet energy demands, especially during fasting periods.

We focus next on the following gap in knowledge in sex differences in metabolism: experiments have shown that increasing hepatic FFA uptake and subsequent FFA oxidation increases gluconeogenesis [50–52]. Yet, women are known to have lower hepatic glucose output than men despite uptaking and oxidizing more FFA [4, 11, 17, 53]. We postulate that sex differences in hepatic glucose output during short-term fasts are influenced, at least in part, by differences in both FFA *and* glycerol, as well as glycogen handling. More precisely, we hypothesize that in the female liver, more lipids are redirected towards carbohydrate metabolism to contribute to glucose production. However, the tendency of the female liver to conserve glycogen hinders a net increase in glucose output.

To test the aforementioned hypothesis, we began by analyzing the upstream fluxes responsible for hepatic glucose output. We note first that in addition to uptaking more FFA, the female liver also uptakes more glycerol than the male liver: 25% and 14% more glycerol uptake in the last 12 hours of a 24-hour fast following high-carbohydrate and high-fat meals, respectively. In a short-term fast, the liver produces glucose through both glycogenolysis and gluconeogenesis. The major substrates of gluconeogenesis are pyruvate (a carbohydrate precursor), glycerol (a fat precursor), and amino acids [59]. Figure 9 shows a comparison between the sexes of the hepatic fluxes involved in glucose production. There is a marginal difference between the sexes in the gluconeogenic flux from pyruvate (*ϕ*_PYR—GAP_). However, for a high-carbohydrate meal, the gluconeogenic flux from glycerol (*ϕ*_GRP—GAP_) is 11% higher in the female model, and glycogenolysis is 7% lower in the female model than in the male model. In the case of a high-fat meal, the gluconeogenic flux from glycerol (*ϕ*_GRP—GAP_) is 9.5% higher in the female model, with glycogenolysis 7% lower compared to the male model. These results suggest that although females take up more FFA during fasting, FFA may be oxidized for energy or re-esterified to TG in the liver. At the same time, glycerol, the other product of lipolysis, is uptaken at higher rates in the female liver and stimulates gluconeogenesis via glycerol oxidation (*ϕ*_GRP—GAP_). But the concurrent decrease in glycogenolysis in females may explain why the net glucose output is lower during fasting. We summarized in Fig 10 the sex-related differences in hepatic metabolic fluxes contributing to the sexual dimorphism in hepatic glucose output during short-term fasts.

**Fig 9.**
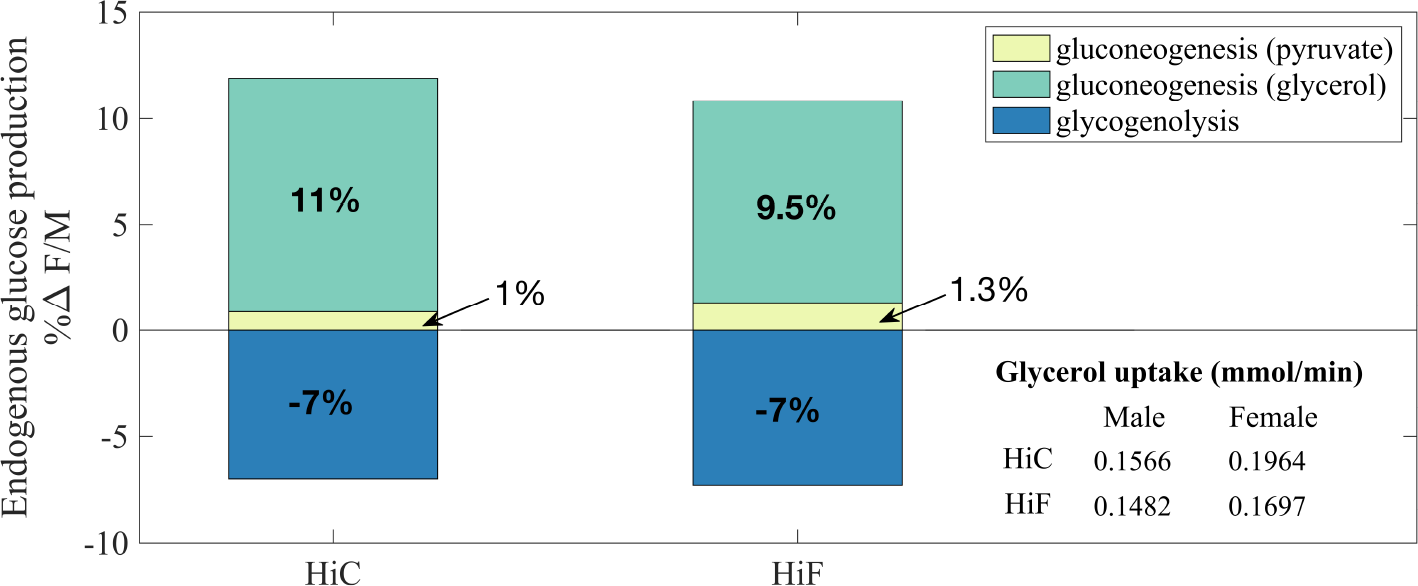
Change in hepatic energy metabolism with fasting. Values represent averages over the last 12 hours of a 24-hour fast following a single 800 kcal meal. HiC, high-carbohydrate meal; HiF, high-fat meal. %Δ F/M refers to the percent relative difference between the sexes. It calculated as (female flux/male flux −1)*×*100.

**Fig 10.**
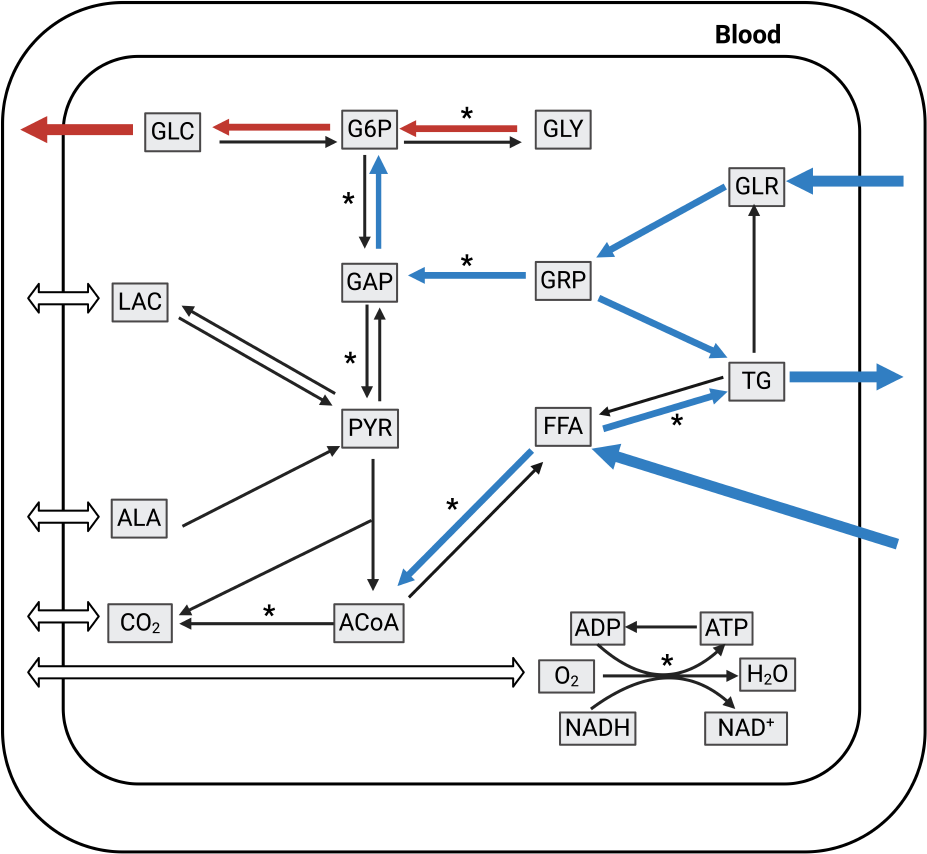
Sex-related differences in liver metabolic pathways during the postabsorptive phase (*>*12 h). Rates higher (lower) in the female model compared to the male model are shown in blue (red). 9 substrates are transported between blood and tissues (open double-sided arrows). Single-sided arrows indicate the direction of transport flux, which varies between the sexes. However, we note that these arrows would be more accurately depicted as double-sided arrows since substrates can be either taken up or released. Pathways marked with an asterisk (*) are composed of multiple reaction steps but grouped together as a single step in this model. Substrate abbreviations are listed in Table 4

Turning to fat metabolism in adipose tissue, our results show that the male model stores more fat as TG in adipose tissue than the female model following a high-fat meal. However, this sex difference in TG accumulation is small and subsides for the case of a high-carbohydrate meal (see Fig 11a). The rate of fat breakdown (lipolysis) in adipose tissue is 22% higher in the female model compared to the male model during the postabsorptive phase (*>*6h; see Fig 11b), despite the two sexes having similar levels of TG accumulation in adipose tissue. This result is consistent with our modeling assumption that body composition influences the extent of regional substrate oxidation: a higher basal body fat percentage in females (29% in the female model vs. 16% in the male model) would result in greater regional lipolysis [18, 28], assuming similar substrate oxidation efficiencies. Notably, Fig 11c shows that greater amounts of TG are released into the circulation by the female liver during the short-term fast irrespective of meal composition. Hepatic fat is mainly delivered to adipose tissue for the purpose of lipolysis. During this time and following a high-carbohydrate meal, skeletal muscle and the GI tract in the female model also uptake more fat than the male model, albeit to a smaller extent than adipose tissue. Following a high-fat meal, skeletal muscle in both models switches from uptake to release of TG, and the male model more so than the female model—an observation consistent with our earlier result that skeletal muscle in females tends to store more IMCL (refer to Fig 7).

**Table 4.**
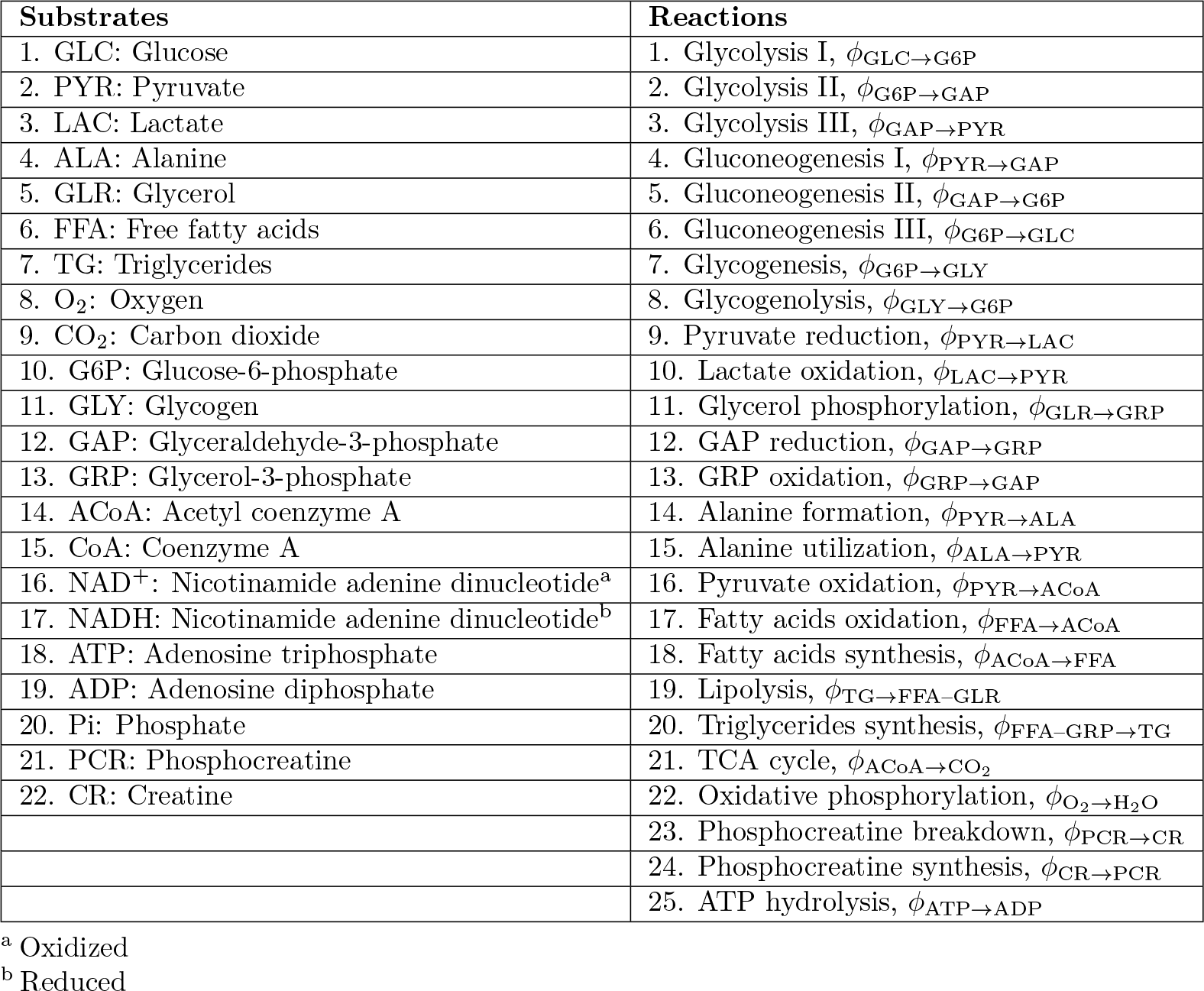
List of substrates and metabolic reactions.

| Substrates | Reactions |
| --- | --- |
| 1. GLC: Glucose | 1. Glycolysis I, $\phi_{\text{GLC} \rightarrow \text{G6P}}$ |
| 2. PYR: Pyruvate | 2. Glycolysis II, $\phi_{\text{G6P} \rightarrow \text{GAP}}$ |
| 3. LAC: Lactate | 3. Glycolysis III, $\phi_{\text{GAP} \rightarrow \text{PYR}}$ |
| 4. ALA: Alanine | 4. Gluconeogenesis I, $\phi_{\text{PYR} \rightarrow \text{GAP}}$ |
| 5. GLR: Glycerol | 5. Gluconeogenesis II, $\phi_{\text{GAP} \rightarrow \text{G6P}}$ |
| 6. FFA: Free fatty acids | 6. Gluconeogenesis III, $\phi_{\text{G6P} \rightarrow \text{GLC}}$ |
| 7. TG: Triglycerides | 7. Glycogenesis, $\phi_{\text{G6P} \rightarrow \text{GLY}}$ |
| 8. O<sub>2</sub>: Oxygen | 8. Glycogenolysis, $\phi_{\text{GLY} \rightarrow \text{G6P}}$ |
| 9. CO<sub>2</sub>: Carbon dioxide | 9. Pyruvate reduction, $\phi_{\text{PYR} \rightarrow \text{LAC}}$ |
| 10. G6P: Glucose-6-phosphate | 10. Lactate oxidation, $\phi_{\text{LAC} \rightarrow \text{PYR}}$ |
| 11. GLY: Glycogen | 11. Glycerol phosphorylation, $\phi_{\text{GLR} \rightarrow \text{GRP}}$ |
| 12. GAP: Glyceraldehyde-3-phosphate | 12. GAP reduction, $\phi_{\text{GAP} \rightarrow \text{GRP}}$ |
| 13. GRP: Glycerol-3-phosphate | 13. GRP oxidation, $\phi_{\text{GRP} \rightarrow \text{GAP}}$ |
| 14. ACoA: Acetyl coenzyme A | 14. Alanine formation, $\phi_{\text{PYR} \rightarrow \text{ALA}}$ |
| 15. CoA: Coenzyme A | 15. Alanine utilization, $\phi_{\text{ALA} \rightarrow \text{PYR}}$ |
| 16. NAD<sup>+</sup>: Nicotinamide adenine dinucleotide<sup>a</sup> | 16. Pyruvate oxidation, $\phi_{\text{PYR} \rightarrow \text{ACoA}}$ |
| 17. NADH: Nicotinamide adenine dinucleotide<sup>b</sup> | 17. Fatty acids oxidation, $\phi_{\text{FFA} \rightarrow \text{ACoA}}$ |
| 18. ATP: Adenosine triphosphate | 18. Fatty acids synthesis, $\phi_{\text{ACoA} \rightarrow \text{FFA}}$ |
| 19. ADP: Adenosine diphosphate | 19. Lipolysis, $\phi_{\text{TG} \rightarrow \text{FFA} - \text{GLR}}$ |
| 20. Pi: Phosphate | 20. Triglycerides synthesis, $\phi_{\text{FFA} - \text{GRP} \rightarrow \text{TG}}$ |
| 21. PCR: Phosphocreatine | 21. TCA cycle, $\phi_{\text{ACoA} \rightarrow \text{CO}_2}$ |
| 22. CR: Creatine | 22. Oxidative phosphorylation, $\phi_{\text{O}_2 \rightarrow \text{H}_2\text{O}}$ |
| | 23. Phosphocreatine breakdown, $\phi_{\text{PCR} \rightarrow \text{CR}}$ |
| | 24. Phosphocreatine synthesis, $\phi_{\text{CR} \rightarrow \text{PCR}}$ |
| | 25. ATP hydrolysis, $\phi_{\text{ATP} \rightarrow \text{ADP}}$ |

<sup>a</sup> Oxidized
<sup>b</sup> Reduced

**Fig 11.**
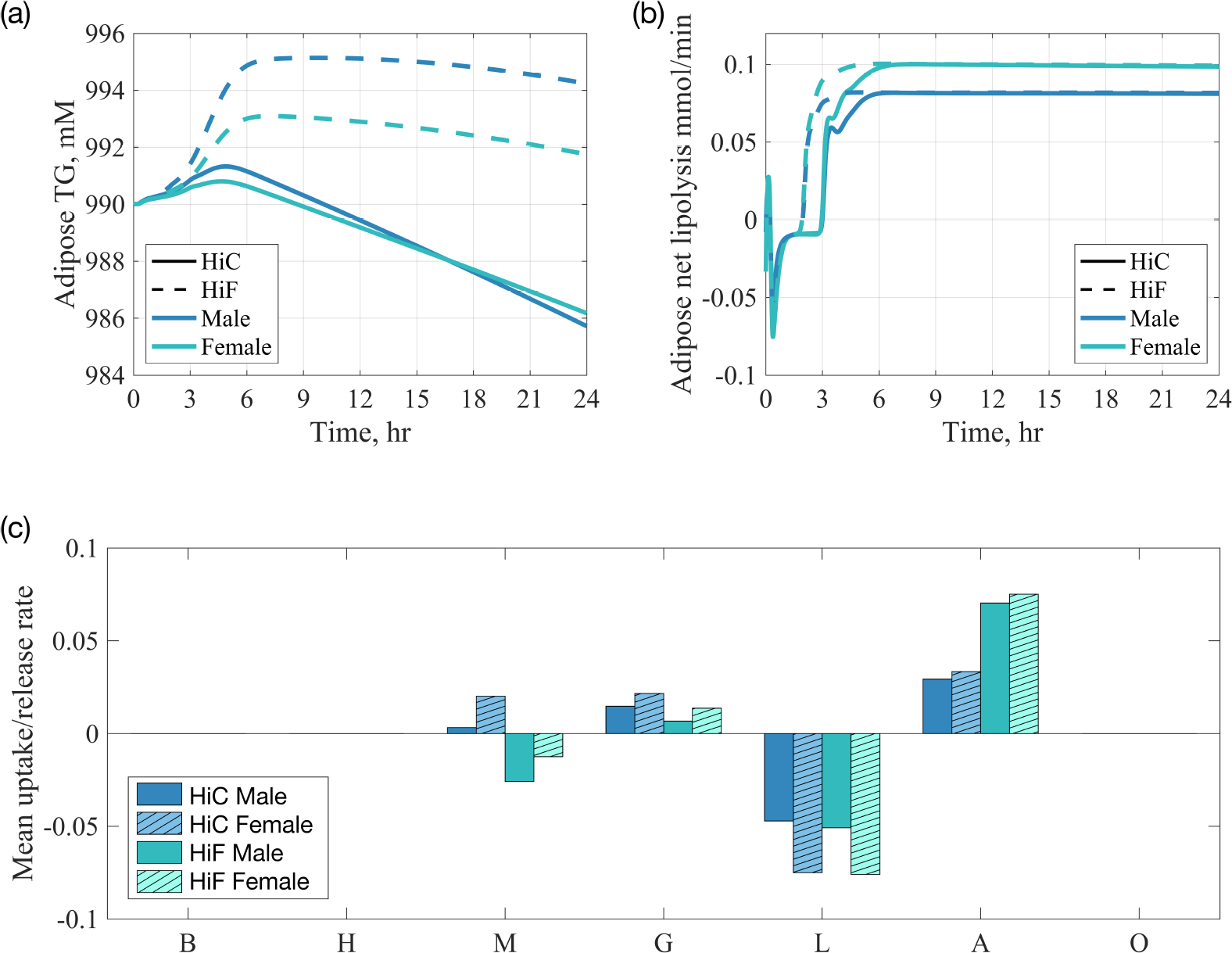
Fat metabolism during a short-term fast (24h) following a single 800 kcal meal. HiC, high-carbohydrate meal; HiF, high-fat meal. (a) Adipose tissue TG concentration; (b) Net lipolysis rate (the difference between TG breakdown and TG synthesis) in adipose tissue ; (c) average uptake (or release) rate (mmol/min) of TG per organ during the last 12 hours of the fasting period following the HiC and HiF meals, respectively; mM, mmol/L. B, brain; H, heart; M, skeletal muscle; G, GI tract; L, liver, A, adipose tissue; O, other tissues.

The exchange of fat, either as TG or FFA, between organs and tissues highlights a metabolic exchange between the liver and adipose tissue. This inter-organ metabolic pathway is more prominent in the female model (Figs 8,9, and 11). Overall the female model has a greater capacity to remove fat from the liver through TG secretion, which is exported to adipose tissue for either storage as TG (e.g., following a high-fat meal) or to be broken down into FFA. For the later, our results show that FFA is then returned to the liver during a short-term fast. This cycle may be the result of a trade-off of efficiencies: adipose tissue is reasonably efficient at extracting FFA from chylomicrons (large triglyceride-rich lipoproteins) while the liver is efficient at converting different gluconeogenic precurssors (e.g., pyruvate, glycerol) into glucose to meet systemic energy needs. Importantly, we note that the hepatic TG pool is not a major energy store for the rest of the body (that function is performed by the TG stored in adipose tissue) but appears to be a local store for hepatic needs, and the stored TG also acts as a substrate for hepatic secretion of fat into the bloodstream (see Fig 10).

#### 2.4.3 Metabolic flux sensitivity to body composition

Our model predicted that sex-related differences in metabolic responses are small during the absorptive phase but become more pronounced during fasting. We argued that the majority of these differences are attributable to differences between sexes in liver and adipose tissue function. Considering the significance of body composition in metabolism, we sought to examine the robustness of our findings in response to variations in body composition. Specifically, we aimed to assess how changes in fat mass and muscle mass impact key metabolic fluxes, such as glycolysis, gluconeogenesis, glycogenesis, glycogenolysis in the liver, and lipolysis in adipose tissue. These fluxes are key in driving whole-body sex differences, as outlined in Table 3. Figures 12 and 13 present the results of a local sensitivity analysis of the mentioned fluxes when fat mass or skeletal muscle mass is varied by either a 5% increase or decrease in both male and female models. Our focus was on the postabsorptive phase, where key sex differences become apparent.

**Fig 12.**
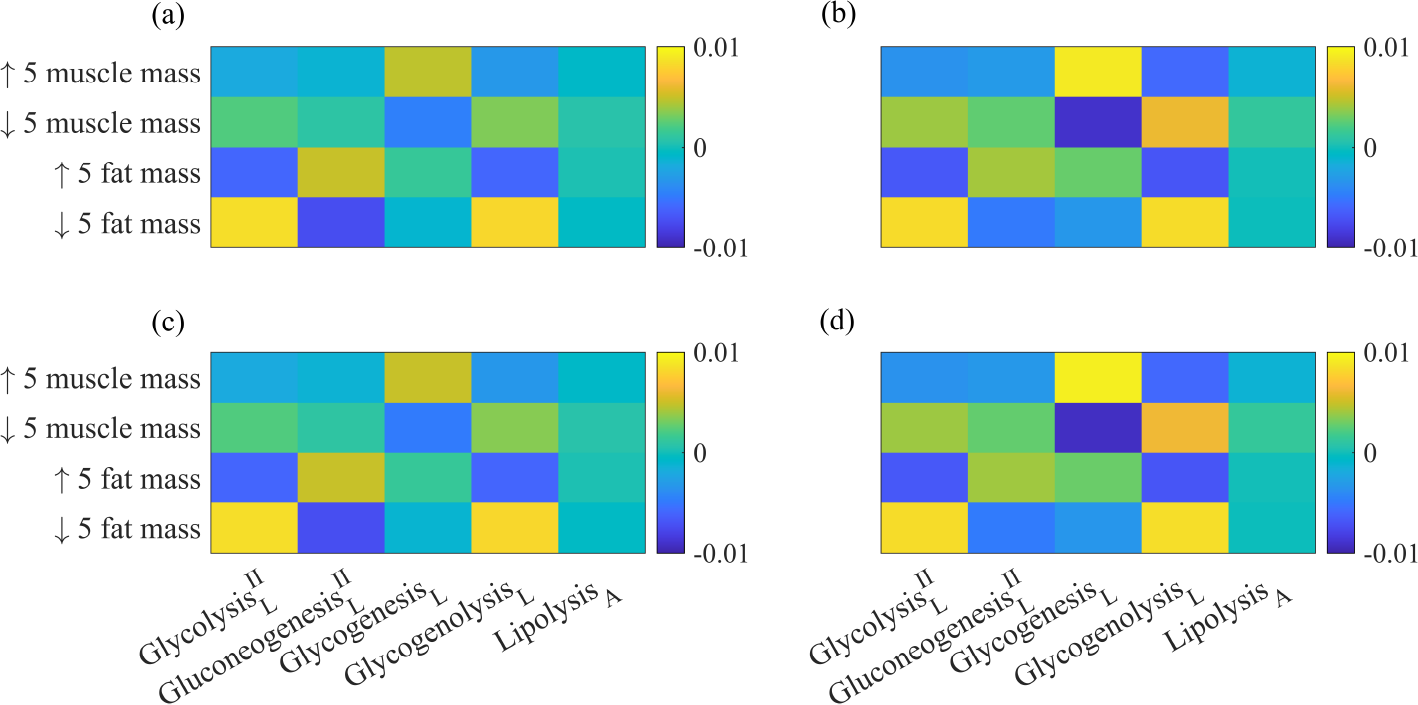
Results of local sensitivity analysis at 9 hours post-meal. (a) Male model and HiC; (b) female model and HiC; (c) male model and HiF; (d) Female model and HiF. HiC, high-carbohydrate meal; HiF, high-fat meal. Glycolysis II, *ϕ*_G6P*→*GAP_; gluconeogenesis II, *ϕ*_GAP*→*G6P_; glycogenesis, *ϕ*_G6P*→*GLY_; glycogenolysis, *ϕ*_GLY*→*G6P_, lipolysis, *ϕ*_TG*→*FFA–GLR_.

**Fig 13.**
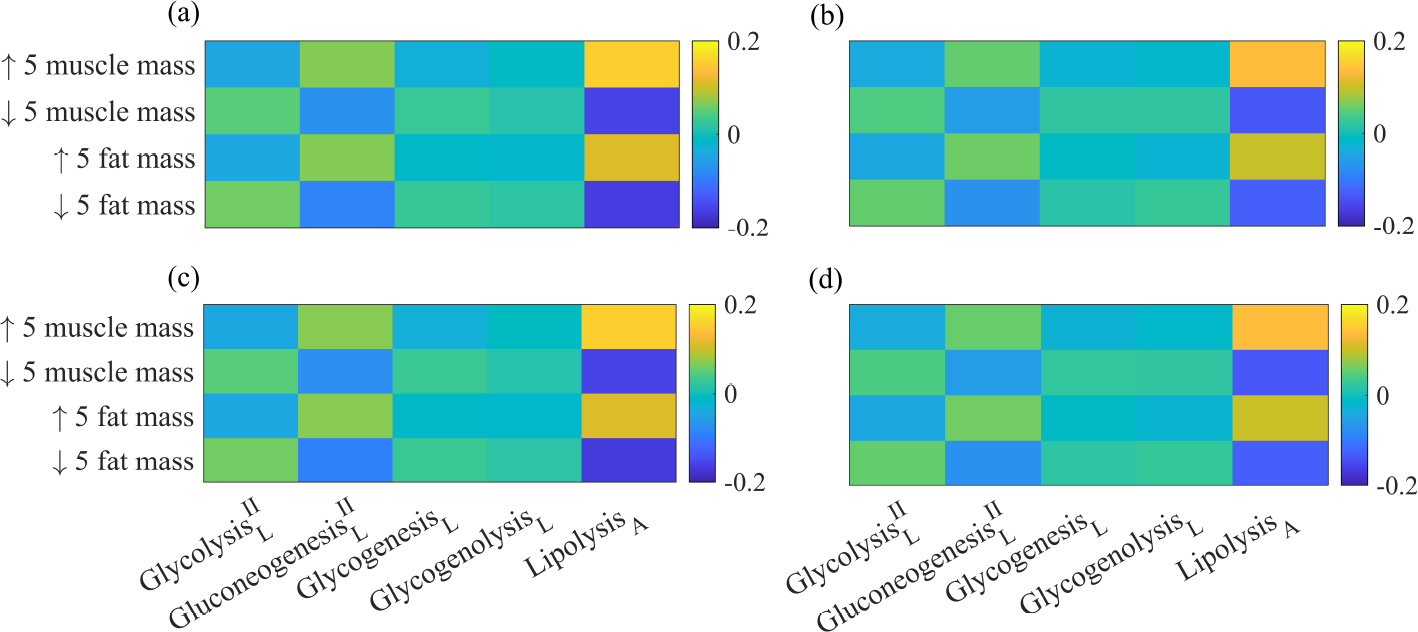
Results of local sensitivity analysis at 24 hours post-meal. (a) Male model and HiC; (b) female model and HiC; (c) male model and HiF; (d) Female model and HiF. HiC, high-carbohydrate meal; HiF, high-fat meal. Glycolysis II, *ϕ*_G6P*→*GAP_; gluconeogenesis II, *ϕ*_GAP*→*G6P_; glycogenesis, *ϕ*_G6P*→*GLY_; glycogenolysis, *ϕ*_GLY*→*G6P_, lipolysis, *ϕ*_TG*→*FFA–GLR_.

Figure 12 shows sensitivity results 9 hours after a meal, while Fig 13 shows the results 24 hours after a meal. In general, at 9 hours post-meal (Fig 12), the male model shows that a 5% increase or decrease in fat mass induces larger changes compared to the same variation in muscle mass in either direction. Across meal types, an increase in fat mass augments gluconeogenesis but diminishes glycolysis and glycogenolysis. The opposite holds for a decrease in fat mass. In the female model, greater variations are observed for changes in muscle mass. A 5% increase in muscle mass results in increased hepatic glycogenesis and reduced hepatic glycogenolysis, with the reverse occurring for a decrease in muscle mass. As the fasting period extends to 24 hours (Fig 13), lipolysis in adipose tissue exhibits more substantial variations than hepatic fluxes, particularly in the male model compared to the female model. During this stage of fasting, changes in fat mass or muscle mass lead to similar sensitivity indices in magnitude, but the direction of change varies. For example, both a decrease in muscle mass and fat mass reduce the lipolysis flux, while the opposite leads to an increase in lipolysis. These results apply to both high-carbohydrate and high-fat diets. Comparing Fig 12 and Fig 13 reveals that hepatic fluxes are most sensitive to body composition during the early postabsorptive period, whereas lipolysis in adipose tissue is most sensitive during the late postabsorptive phase.

Overall, the magnitudes of the sensitivity indices remain small, with *−*0.2 *< S*_*i*_ *<* 0.2 and where *i* represents either fat mass or muscle mass. This observation applies to both models and across both meal types. Such findings provide increased confidence that the model predictions reflect genuine sex-related metabolic differences, regardless of variations in body composition. We also point out that the sensitivity indices at 24 hours post-meal are higher than those at 9 hours post-meal by an order of magnitude. This suggests that the model is more sensitive to changes in body compositions during the latter stages of the postabsorptive period, indicating that sex differences associated with body composition may become more pronounced during longer fasts. This aligns well with our findings that sex differences become more apparent during the late postabsorptive phase (see Table 3). More detail about our sensitivity analysis method and actual values of sensitivity indices are available in S1 Supporting Information (Section 2).

## 3 Discussion

### Variations in whole-body metabolism resulting from dietary factors

Throughout the absorptive phase (0-6h), both sexes experience a marked increase in RQ (*≥* 0.88), indicative of a preference for carbohydrate oxidation. Figure 5 shows the whole-body RQ results for male and female models. During the transition to the postabsorptive phase, RQ values, although decreasing, remain *>* 0.8, suggesting sustained carbohydrate oxidation in the early postabsorptive period (6-12h). This trend is consistent across sexes, showing no significant differences in RQ values. A recent 2021 experimental study [72] also showed no sex-related difference in RQ in lean, healthy young males and females in the short term following a high-fat diet [72].

Importantly, meal composition, rather than sex, exerts the most significant influence on the evolution of whole-body RQ after a single mixed meal. Our results indicate that RQ peaks over an hour earlier for a high-fat meal compared to a high-carbohydrate meal (Fig 6). In line with this observation, the oxidation fractions of carbohydrates, reflecting the extent to which carbohydrates contribute to ATP energy production, peak earlier and remain elevated for 2.5 hours following a high-fat meal compared to a high-carbohydrate meal. Yet, the increased availability of carbohydrate following a high-carbohydrate meal leads to a swift increase in carbohydrate oxidation, surpassing that induced by a high-fat meal after 3 hours.

This seemingly counterintuitive result, considering that a high-carbohydrate meal has 40% more carbohydrate content than a high-fat meal in our simulations, raises questions about the underlying metabolic pathways and energy utilization during the digestion of carbohydrates and fats. Mechanistically for a high-fat meal, the initial spike in RQ can be attributed to the body’s preference for utilizing carbohydrates for quick energy. However, as the digestion and absorption of dietary fat progress, the body gradually shifts to using fatty acids for energy. This transition from carbohydrate to fat metabolism contributes to the eventual decline in RQ. Conversely, with a high-carbohydrate meal, the body promptly responds by prioritizing the export of carbohydrates from the systemic circulation. This is followed by the breakdown and storage of excess carbohydrates as glycogen, primarily in organs like skeletal muscle and the liver. This rapid uptake into organs prevents postprandial hyperglycemia, characterized by elevated blood glucose levels persisting for more than 1-2 hours after food ingestion [61].

The delayed increase in RQ following a high-carbohydrate meal can be attributed to this differential metabolic response: aggressively storing glucose to restore normal glycemia, thus continuing to burn fat to some extent during the initial phase following a high-carbohydrate meal. Then, the body eventually transitions to using carbohydrate as fuel. In summary, the sequence of RQ peaks reflects the dynamic interplay between carbohydrate and fat metabolism during the absorptive phase, illustrating the body’s adaptive mechanisms to efficiently extract energy from different macronutrients over time.

### Variations in whole-body metabolism resulting from sex differences

We observed sex-related differences in whole-body metabolic fluxes that persist across meal types, with more pronounced effects following a high-carbohydrate meal (Table 3). In the absorptive phase, our findings indicate that the female model tends to preserve significantly more fat than the male model. This is evident in the net rate of TG synthesis, which is 21% and 26% higher in the female model for high-carbohydrate and high-fat meals, respectively. Additionally, *de novo* lipogenesis, the process synthesizing FFA from substrates such as glucose and amino acids, is higher in the female model (Table 3). This implies that more glucose is stored as fat in the female model than in the male model during the absorptive phase. Sex differences in glucose metabolism during the absorptive phase are also apparent and suggest a preference in the female model to utilize more carbohydrates to meet energy needs (Table 3). Notably, rates of glycogen breakdown and gluconeogenesis are higher in the female model during this period. Experimental data indicate a complex role of sex hormones, with sexual dimorphisms depending on the individual’s metabolic status. For example, women in the prandial and postprandial states utilize more carbohydrates than men to meet energy needs, while during fasting or exercise, women rely more on fat oxidation than men [6, 7, 10, 13, 62]. Our results align with the former during the absorptive phase, and we will revisit the latter below for the postabsorptive phase.

In the postabsorptive phase, sex differences become more evident in both fat and glucose metabolism and following a high-carbohydrate meal. The female model produces less glucose but oxidizes more fat. Specifically, our female model exhibits a 10% higher *net rate* of glycogenolysis and a 15% lower *net rate* of gluconeogenesis compared to the male model. The female model shows lower rates of glucose production as the decrease in gluconeogenesis dominates. Experimental studies have also demonstrated this sexual dimorphism in glucose production [6, 7]. Additionally, lipolysis is 8% higher in the female model, suggesting a higher reliance on FFA by other organs and tissues during the postabsorptive phase. Following mass action principles, higher glycogen levels may lead to increased rates of glycogen breakdown, and higher fat levels may lead to increased rates of fat hydrolysis. As expected, these rates are higher for the female model, given that the female liver stores more glycogen during the absorptive phase, and the female model has a higher initial body fat percentage (and mass). Mechanistically, the reduced rate of gluconeogenesis in the female model is a consequence of increased systemic lipolysis. Since the rate of glucose output from the liver to the circulation depends on the rate of glucose uptake by other organs, the inherent net rate of gluconeogenesis is reduced in the female model as other organs and tissues supplement their energy needs by uptaking and oxidizing FFAs during the postabsorptive phase. Our predictions aligns with experimental observations indicating that women rely more on fat oxidation than men during fasting [6, 7, 10, 62]. After a high-fat meal, the female model continued to show elevated rates of lipolysis. In general, disparities related to sex are more noticeable in the postabsorptive phase compared to the absorptive phase, particularly following a high-carbohydrate meal compared to a high-fat meal.

### Sex-differences at the organ and tissue levels drive systemic differences

Our analysis revealed sex-related differences during both the absorptive and postabsorptive phases, which persisted for both high-carbohydrate and high-fat meals. Notably, these differences tended to be more pronounced following a high-carbohydrate meal. In the absorptive period, two substrates, namely glycogen and TG, exhibited a significant difference between the sexes (Fig 7). The male model tended to store less glycogen in the liver and skeletal muscle than the female model, consequently producing and oxidizing more glucose for energy. Regarding TG, our models showed similar amounts stored in the liver, while the male model stored significantly less TG in skeletal muscle fibers (approximately 60% less). This aligns with the findings of Devries et al. [3] and Tarnopolsky et al. [8], indicating that men have lower IMCL density compared to women (28%–84% less).

Sex-related differences became more evident during the postabsorptive period (more than 6 hours after a meal). The female liver’s inclination to preserve more glycogen than the male model extended into the postabsorptive phase (Fig 8c). Consequently, the female liver released less glucose into the circulation during that time (Fig 8c). To offset the reduced glucose output, the liver and other organs increased their reliance on FFA oxidation for energy. For example, the female liver took up significantly more FFA to meet its energy needs, and some of that FFA was also re-esterified and released into the bloodstream as TG (Fig 11c). The rate of FFA uptake is directly linked to the substrate’s availability in the circulation [59]. The increased FFA availability is attributed to lipolysis in adipose tissue, the GI tract, and skeletal muscle (Fig 8a). We note that the basal rate of lipolysis in adipose tissue is higher in our female model by 20%, in line with findings from experimental studies [18, 28]. Overall, FFA release is higher from adipose tissue and skeletal muscle in the female model. For the latter, the greater amount of IMCL in the female model explains the higher rate of muscle lipolysis. We identified a candidate mechanism, driven by adipose tissue and the liver, which could mechanistically explain the observed sex differences in hepatic glucose output and FFA availability/oxidation between the sexes. We discuss it next.

### Liver and adipose tissue emerge as hubs of sex-related metabolic differences

Especially during the late postabsorptive phase (more than 12 hours after a meal), our models have highlighted a metabolic exchange between the liver and adipose tissue that enables the body to meet energy requirements when glycogen stores are low or depleted (Fog 10). For instance, we observed that in addition to FFA, the female liver also takes up more glycerol, the other product of lipolysis. On the one hand, when FFA enters hepatic cells, some of it is oxidized for energy, and some is re-esterified to TG. Our models show that these processes occur at higher rates in the female model compared to the male model. On the other hand, the glycerol entering hepatic cells is diverted to promote gluconeogenesis. We observed no sex difference in the rate of gluconeogenesis from pyruvate. In net, the rate of gluconeogenesis is higher in the female model than the male model. Simultaneously, the rate of glycogenolysis is reduced in the female liver to the extent that the net rate of glucose production and subsequent glucose output is lower in the female model compared to the male model. We recall that glycogenolysis and gluconeogenesis contribute almost equally during the postabsorptive phase to glucose production. As such, other organs compensate for glucose by taking up and oxidizing FFA (except the brain).

The TG released by the liver is then taken up by adipose tissue, broken down into FFA, and released into the circulation for re-uptake by liver. This cycle implies that females both synthesize and burn more fat than males during a short-term fast. A result supported by the higher levels of plasma FFA and glycerol during the postabsorptive period and regardless of meal composition (Figs 4c and d4). The greater glycerol rate of appearance is an index of the whole-body lipolytic rate [6]. This metabolic exchange between the liver and adipose tissue is independent of sex, but the difference in body composition and reaction rates between the sexes causes the female model to rely more heavily on it.

Our candidate mechanism could partly explain a current gap in the literature. Increased uptake of FFA is known to increase gluconeogenesis [50–52], but females, who significantly uptake more FFA than males, also produce less glucose [4, 11, 17, 53]. This suggests that an alternate mechanism hinders the increase in gluconeogenesis from leading to increased glucose output. In summary, we posit that gluconeogenesis is increased in females due to the concurrent uptake of FFA *and* glycerol. Glycerol specifically increases gluconeogenesis in the liver, but the reduction in the rate of glycogenolysis is such that the net contribution to glucose output is negative in females. At the core of these sex differences are the liver, which preserves more glycogen in females, and adipose tissue, which provides FFA and glycerol to meet the body’s energy needs. The latter is accentuated in females, who have a higher basal rate of lipolysis regardless of the fact that they have a higher percentage of body fat [18, 28]. Our candidate mechanism is summarized in Fig 10.

## 4 Conclusion

We developed a whole-body, multi-scale, and sex-specific model of energy metabolism. We incorporated metabolic adjustments that connect cellular metabolism in organs to systemic responses following different mixed meals. Our model aligns well with experimental data. According to our predictions, sex-related metabolic differences are more noticeable following a high-carbohydrate meal compared to a high-fat meal, with differences becoming more pronounced during a short-term fast. In summary, women tend to preserve more fat than men during the absorptive period but oxidize significantly more fat during the postabsorptive period. We hypothesized that the increased reliance on fat metabolism in females is influenced by sex differences in the liver and adipose tissue, and we have outlined a candidate underlying mechanism for this sexual dimorphism. Specifically, the female liver conserves more glycogen, leading to reduced glucose output. This decrease in arterial glucose promotes FFA oxidation by other organs and tissues, except the brain. Computational biology offer promising avenues for refining whole-body metabolic models. Integrating sex-specific data and parameters into multi-scale frameworks holds the potential to enhance our understanding of human metabolism and its modulation by sex. By accounting for the intricate interplay between organs, hormones, and metabolic pathways, these models can provide valuable insights into the underlying mechanisms driving sex-specific metabolic responses.

## Appendix A Models

In this section we formulate a mathematical model describing human metabolism and the dynamics of metabolic flexibility. We model seven tissue compartments: brain, heart, liver, GI tract, skeletal muscle, adipose tissue, and “other tissues” (Fig 1). The latter includes the kidneys, upper extremity muscles, and remaining tissues. Each compartment is characterized by dynamic mass balance equations governing 25 cellular metabolic reactions involving 22 substrates (refer to Table 4). All compartments are interconnected via blood circulation. This model represents an extension of our previously published whole-body model [20] that focused on fuel homeostasis during exercise without food intake. Our model accounts for the synthesis and transport of each substrate across compartments.

Our focus here is to investigate the temporal evolution of substrate concentrations following a single meal and short fast of less than 24 hours. A meal is defined as a combination of carbohydrates and fat. It is important to note that protein is not included in our modeling. The investigation involves two meal compositions: a high-carbohydrate meal, with 90% of caloric intake from carbohydrates and 10% from fat; and a high-fat meal, with an equal distribution of 50% caloric intake from carbohydrates and 50% from fat. Carbohydrate specifically refers to glucose, and fat refers to TG. Table 2 provides details on meal composition. We assume the usage of glucose and TG by all tissues and organs to be dependent on the amount of each substrate in the blood plasma. We construct the model one compartment at a time and start with the blood compartment as this acts as the main transport medium for substrates between the different organs and tissues of the body. We note that no synthesis of glucose or TG occurs in the circulation.

### A.1 Blood compartment and dietary intake

The model simulates a meal by adding glucose and TG into the bloodstream. The model blood compartment serves as the primary conduit for substrate transport among the seven tissue compartments. Besides glucose and TG, the transport of the following substrates also occurs between blood and tissue: pyruvate, lactate, alanine, glycerol, free fatty acids, oxygen, and carbon dioxide. The alteration in the concentration of a substrate in the circulation is governed by the cumulative transport fluxes among all organs and the circulation, and, in the case of glucose and TG, a source term.

Consequently, the plasma concentration of a substrate *i* is modified as follows:

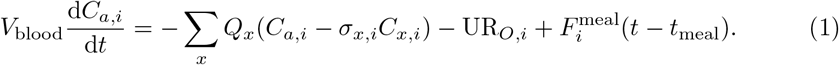

Here, *V*_blood_ is the blood volume, *Q*_*x*_ is the blood flow to/from organ *x, C*_*a,i*_ is the plasma concentration of substrate *i, C*_*x,i*_ is the corresponding concentration of substrate *i* in organ *x*, and *σ*_*x,i*_ is the partition coefficient of substrate *i*. These partition coefficients, unchanging, indicate the relative distribution of metabolites between blood and organs at rest. The expression *Q*_*x*_(*C*_*a,i*_ *− σ*_*x,i*_*C*_*x,i*_) characterizes the net rate of substrate absorption into organ *x*, with the negative sign indicating transport out of the circulation. *x* represents brain, heart, skeletal muscle, GI tract, liver, and adipose tissue. The “other tissues” compartment includes the kidneys, upper extremity muscles, and remaining tissues, and lacks any metabolic reactions. It functions as a source or sink for the nine substrates exchanged between blood and tissues. In the case of other species found exclusively in tissues, the source or sink terms are nil. Let UR_*O,i*_ denote the source or sink term for substrate *i* in tissue *O*, which represents *other tissues*. A positive value indicates uptake of the corresponding substrate, while a negative value indicates release into the circulation. The source or sink values are set to maintain the overall mass balance of the whole-body model at rest. However, for substrates such as lactate and glycerol, which can be excreted by the kidneys when present in excess in the circulation, the source or sink terms are dynamic and can model excretion. When *I* represents lactate or glycerol,

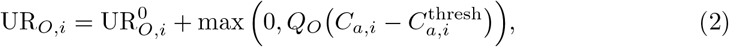

where 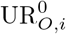 is the basal rate of uptake or release of substrate *i* from *other tissues, Q*_*O*_ is the blood flow to *other tissues, C*_*a,i*_ is the blood concentration of substrate *i*, and 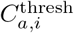 is the corresponding excretion threshold. The kidney is both a lactate producer and consumer. For instance, in the absorptive state, kidneys act as recipients of the lactate shuttle, and when in excess, they serve as sites of net lactate disposal [73, 74]. Given that normal lactate levels are less than 2 mM [75], 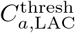 is set to 2 mM. According to Nelson et al. [76], plasma glycerol concentrations above 0.327 *±* 0.190 mM are associated with urinary glycerol excretion. Thus, 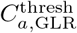 is set to 0.33 mM. For other substrates—glucose, pyruvate, alanine, TG, oxygen, and carbon dioxide, 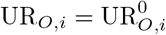 .

We model the rate of appearance of glucose and TG in the blood using *modified* equations introduced by Pearson et al. [41]:

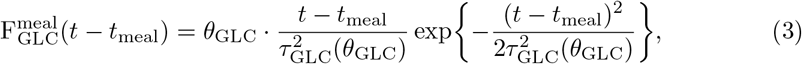

and

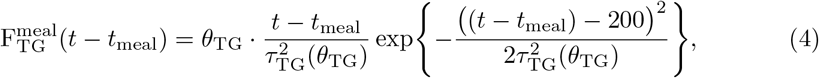

where *t* and *t*_meal_ indicate time (in min), and *t*_meal_ specifically marks the beginning of a meal. According to Frayn [59], glucose peaks 30-60 mins after a meal, and TG peaks 3-4 hours after a meal. The term (*t − t*_meal_ *−* 200) models this delayed peak in TG appearance. For *i* ≠ GLC or TG, 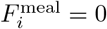. The functions *θ*_GLC_ and *θ*_TG_ are the concentrations of exogenous glucose and TG, respectively, and determine the magnitude of substrate excursion entering the circulation. The functions *τ*_GLC_(*θ*_GLC_) and *τ*_TG_(*θ*_TG_) determine the timescale of substrate release into the blood. While Pearson et al. [41] defined *τ*_GLC_ and *τ*_GLC_ as constants, we define them as dynamic parameters. We model *τ*_GLC_(*θ*_GLC_) and *τ*_TG_(*θ*_TG_) as sigmoids, which depend on the size of the ingested meal, and lead to the desired peak times of release:

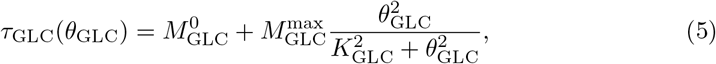

and

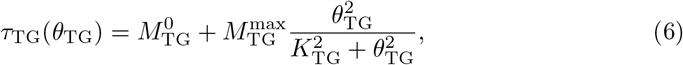

where 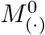 (min) is fixed and represents the basal gastric emptying time for an exogenous substrate, 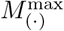 (min) is the maximal gastric emptying time, and *K*_(*·*)_ (mmol) is the half-emptying threshold in the gut. Our modification leads to a more physiological representation of substrate release into the peripheral circulation following the intake of a mixed meal compared to the initial model presented in Ref. [41]. In all of the above equations, *t, τ*_GLC_, and *τ*_TG_ are expressed in minutes. We set 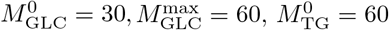, and 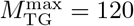, in accordance with Ref. [59].

### A.2 Pancreatic hormones

The pancreas has a dual role in regulating macronutrient absorption and metabolism. It secretes digestive enzymes (exocrine function) and pancreatic hormones (endocrine function) [59]. Acinar cells, also known as exocrine cells, produce pancreatic juices containing digestive enzymes like amylase, pancreatic lipase, and trypsinogen. Pancreatic hormones, on the other hand, are released in an endocrine manner—directly into the bloodstream. Endocrine cells aggregate to form the islets of Langerhans, making up only 1–2% of the entire organ. These islets resemble small, island-like structures within the exocrine pancreatic tissue [77]. The endocrine pancreas is a key contributor to blood glucose regulation by producing the hormones insulin and glucagon. While glucagon increases blood glucose levels, insulin acts to decrease them. Specialized cells, namely the glucagon-producing *α*-cells (15–20% of total islet cells) and insulin-producing *β*-cells (65–80% of total cells), are responsible for synthesizing these hormones [77]. In this model, there is no distinct pancreatic compartment; instead, the release and elimination of hormones from the plasma are described through phenomenological functions that capture the established relationships between plasma glucose levels and hormone concentrations (Fig 14).

**Fig 14.**
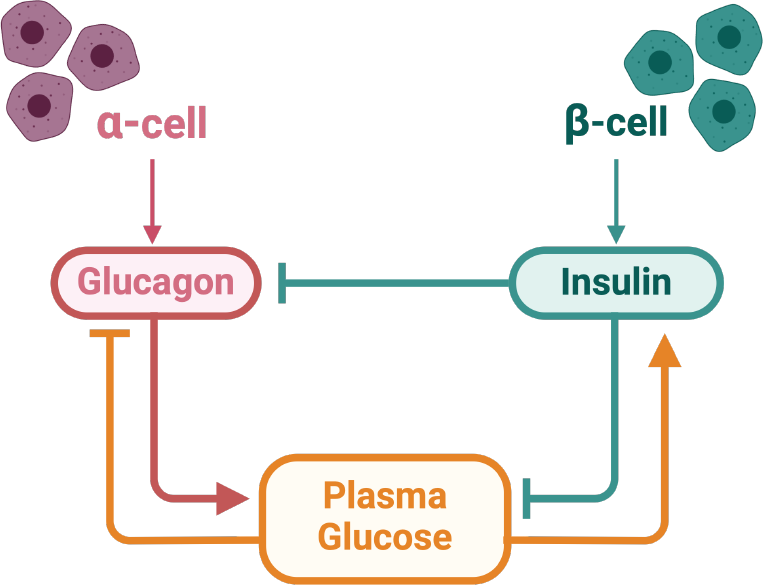
Endocrine control of plasma glucose. Blunt arrows indicate inhibition, while pointed arrows indicate stimulation. Regarding the effects of insulin and glucagon on plasma glucose, arrows denote the signaling of glucose utilization (blunted arrows) or production (pointed arrows) by other organs.

Equations (7) and (8) below describe the dynamics of insulin and glucagon, respectively; together, they represent the pancreatic hormone responses.

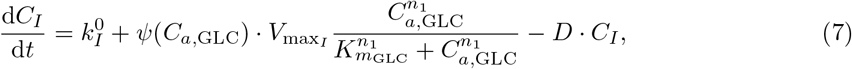

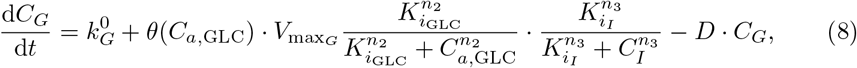

with

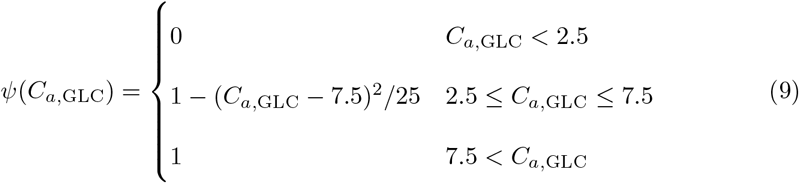

and

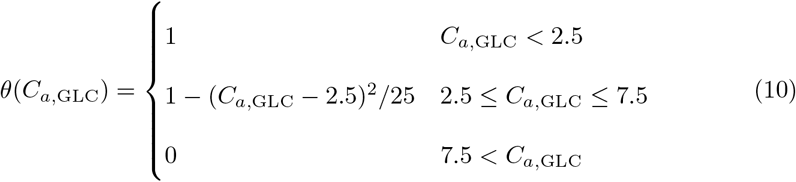

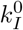 and 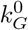 represent the basal rates of insulin and glucagon secretion, respectively, while *V*_max,I_ and *V*_max,G_ represent the maximum rate coefficients. *K*_*m*,GLC_ is the glucose concentration at which half of *V*_max,I_ is achieved. Similarly, *K*_*i*,GLC_ and *K*_*i,I*_ serve as inflection points for the glucose and insulin sigmoids, respectively. These values, known as inhibition constants, indicate the concentration required for half-maximum inhibition. The exponents *n*_1_, *n*_2_, and *n*_3_ denote the slopes of the sigmoids, characterizing sensitivity of responses. *D* represents the disappearance rate of insulin and glucagon. The parameter values are sourced from literature or determined through fitting to experimental data (refer to Section B). Specifically, *K*_*m*,GLC_ = 7 mM [59, 78], *K*_*i*,GLC_ = 20 mM [79], *K*_*i,I*_ = 90 pM [80], *n*_1_ = 7 [81], and *D* = 0.1 min^*−*1^ [82]. The remaining parameters—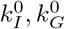, *V*_max,I_, *V*_max,G_, *n*_2_, and *n*_3_—are estimated from the available data [54, 55, 58].

These responses (Eqs (7) and (8)) are sigmoid functions that range between basal hormone secretion rates (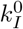 and 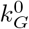) and maximal hormone secretion rates (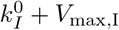 and 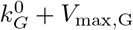). They monotonically increase with rising blood glucose levels for insulin (Eq (7)) and decrease for glucagon (Eq (8)). In terms of regulation, glucose stimulates insulin release while suppressing glucagon secretion [59]. Furthermore, recent work by Vergari et al. [80] indicates direct paracrine inhibition of glucagon secretion by insulin. The relationships between glucose and pancreatic hormones follow characteristic sigmoid dose–response curves [32, 59], which we modeled by a Michaelis-Menten-like hyperbolic functions. We assumed that the hormones degrade at a rate linearly proportional to their respective concentrations (*D · C*_*I*_ and *D · C*_*G*_). We introduced the terms *θ*(*C*_*a*,GLC_) and *ψ*(*C*_*a*,GLC_) to capture the influence of glucose on maximum rate coefficients. Specifically, the *V*_max_ values for hormone secretion correlate with the prevailing glucose concentration [83]—*ψ*(*C*_*a*,GLC_) increases with glucose, while *θ*(*C*_*a*,GLC_) decreases. Both are formulated to maintain a steady-state plasma glucose concentration of 5 mM [84].

### A.3 Organ and tissue compartments

In the whole-body model, a spatially lumped capillary blood domain facilitates the exchange of nutrients and metabolic substrates with spatially lumped domains of tissue cells, each representing one of the modeled organs or tissues. Each organ compartment, except “other tissues”, is characterized by a set of organ-specific metabolic reactions. Figure 15 provides a comprehensive overview of all the metabolic pathways represented in our models, and Table 5 delineates tissue-specific metabolic pathways.

**Table 5.**
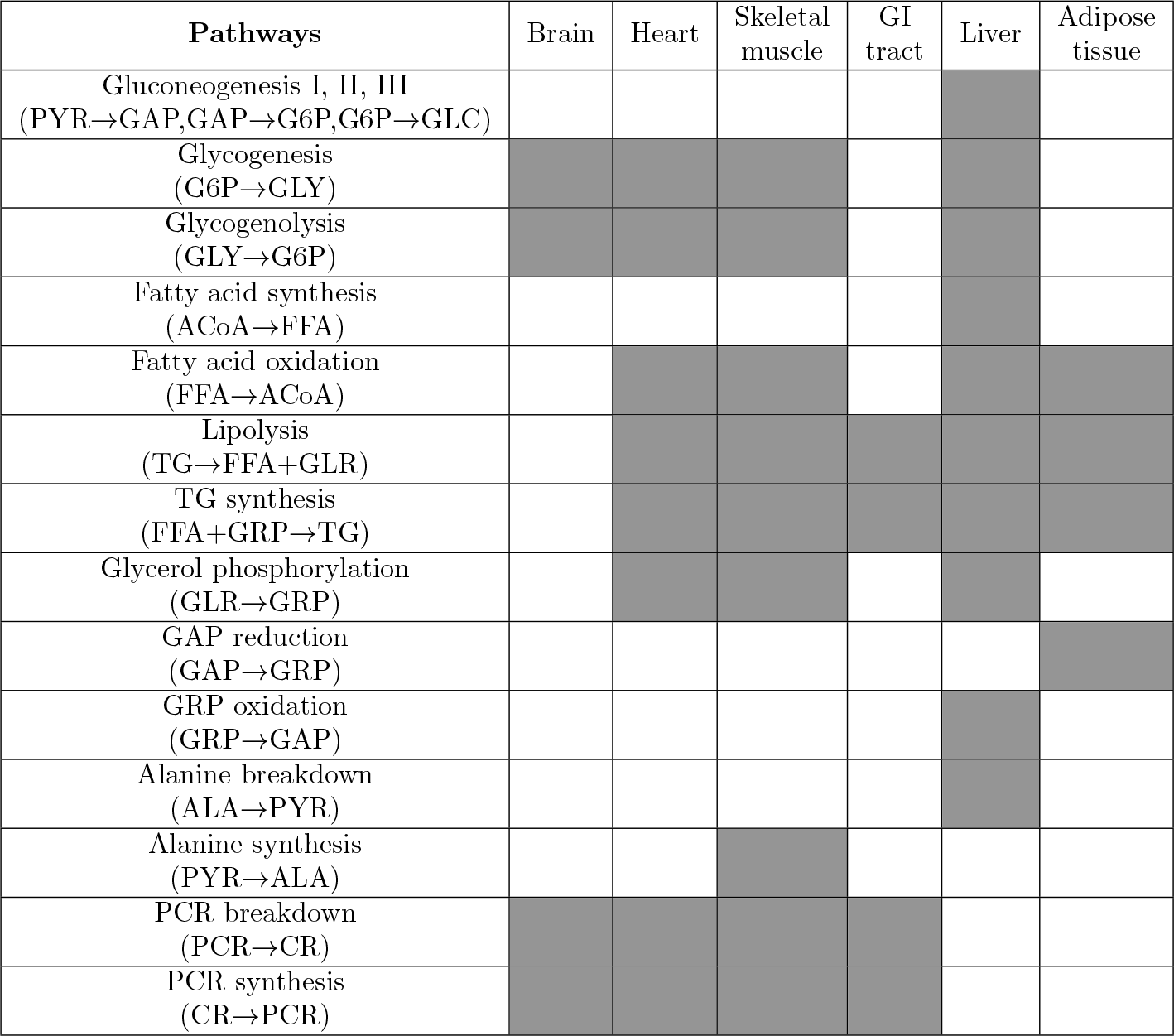
Map of tissue-specific metabolic pathways. A filled box means the existence of the corresponding pathway. In addition to the common pathways depicted in Fig. 15, each tissue has its own different set of metabolic pathways. Table reproduced from Ref. [20] with permission.

**Fig 15.**
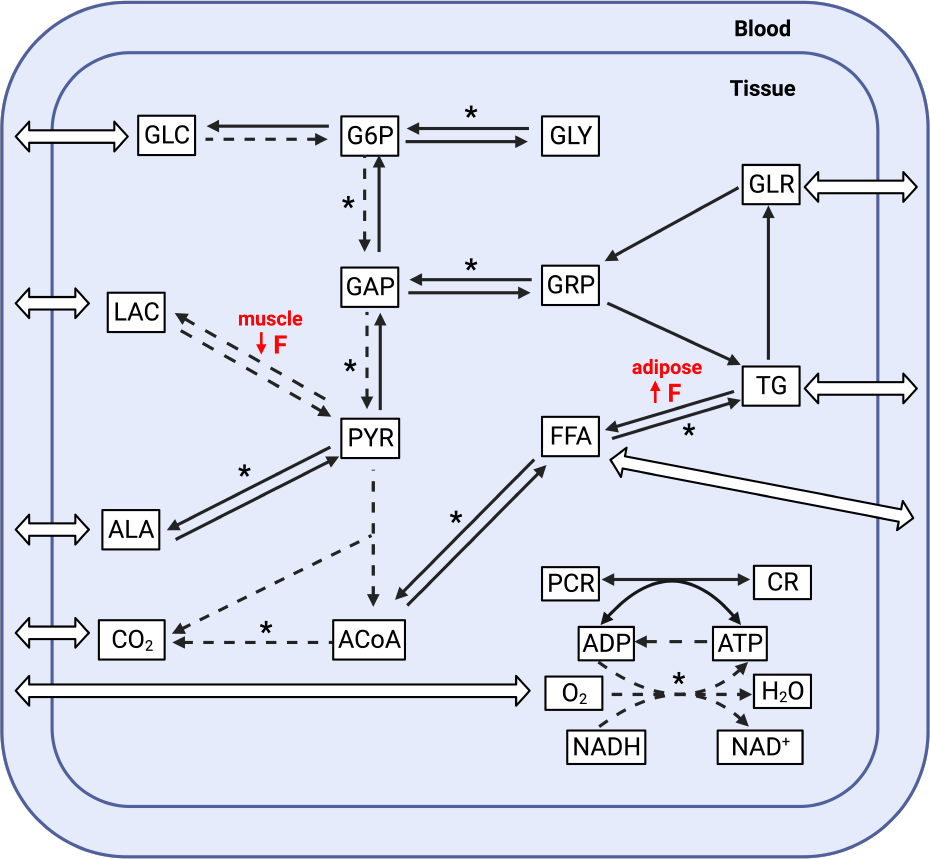
Union map of all-organ metabolic pathways. 9 substrates are transported between blood and tissues (open arrows). Black arrows are tissue-specific pathways, whereas dashed arrows represent common pathways found in all tissues. Pathways marked with an asterisk (*) are composed of multiple reaction steps but grouped together as a single step in this model. Reaction rates in females that are significantly different from males at rest are marked by an arrow indicating the direction of change and the symbol F. Substrate abbreviations are listed in Table 4. Figure reproduced from Abo et al. [20] with permission.

The general form of the mass balance equation for chemical species *i* in tissue *x* is

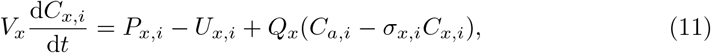

where *V*_*x*_ is a constant parameter representing the volume of tissue *x*. The terms *P*_*x,i*_ and *U*_*x,i*_ denote the production and utilization rates of substrate *i* in tissue *x*, respectively. *Q*_*x*_ is fixed and corresponds to the blood flow to tissue *x*, while *C*_*a,i*_ and *C*_*x,i*_ represent the arterial and tissue concentrations of substrate *i* respectively. The parameter *σ*_*x,i*_ denotes the partition coefficient of substrate *i*, which remains fixed and reflects the relative distribution of metabolites between blood and tissues at rest. The first two terms on the right side of Eq (11) express the net metabolic reaction rate of substrate *i* in tissue *x*. The third term, *Q*_*x*_(*C*_*a,i*_ *− σ*_*x,i*_*C*_*x,i*_), is the net rate of absorption or release of substrate *i* in tissue *x*. It is worth noting that only 9 substrates (substrates 1 to 9 in Table 4) are transported between blood and tissue.

In the absorptive period following a meal rich in carbohydrates or fats, hyperglycemia and/or hypertriglyceridemia may occur. Hyperglycemia is typically defined by blood glucose levels exceeding 6.1 mM [78], while hypertriglyceridemia is indicated by blood TG levels surpassing 2 mM [59]. Hyperglycemia plays a role in stimulating glucose uptake by splanchnic (liver and GI tract) and peripheral (mainly skeletal muscle) tissues. Liver cells are primarily equipped with the GLUT-2 glucose transporter, which does not respond to insulin. This implies that the rate and direction of glucose movement across the hepatocyte membrane depend on the relative concentrations of glucose inside and outside the cell. In skeletal muscle, glucose uptake is predominantly mediated by the insulin-sensitive glucose transporter, GLUT4. The conventional expectation is that hyperinsulinemia follows hyperglycemia, leading to the stimulation of glucose uptake. This occurs through an increase in the number of GLUT4 transporters at the cell membrane and enhanced disposal of glucose within the cell [59]. However, it is noteworthy that hyperglycemia, even in the absence of hyperinsulinemia, independently stimulates muscle glucose uptake [85]. To enhance the model’s ability for maintaining glucose homeostasis under different metabolic conditions, we assume that glucose uptake into the liver and skeletal muscle is stimulated by hyperglycemia, regardless of whether it is coupled with hyperinsulinemia or not. In these organs, the partition coefficient for glucose, which modulates the organ’s capacity for uptake or release of glucose, is a dynamic parameter

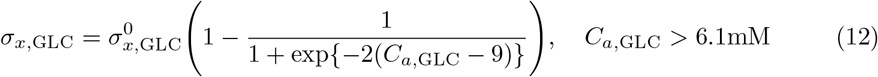

and 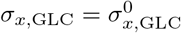, otherwise. In this equation, *x* denotes the liver or skeletal muscle, and 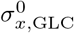 represents the partition coefficient under normal glucose levels. The slope of the sigmoid is set to 2 to simulate the rapid clearance of glucose during hyperglycemia, in line with expectations for healthy individuals [59, 78]. The inflection point of the sigmoid is chosen as 9 so that glucose levels above 6.1 mM start to elicit a response. Similarly, in the fed state, adipose tissue serves as the primary site for the uptake of dietary TG. Therefore, during postabsorptive hypertriglyceridemia, we assume that the influx of TG into adipose tissue is modulated as follows:

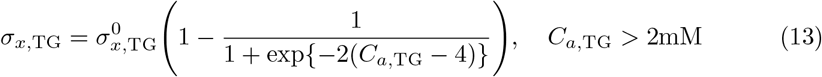

where *x* represents adipose tissue, and 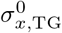 denotes the partition coefficient under normal TG levels. In this case as well, the slope of the sigmoid is set to 2, and the inflection point is chosen so that TG levels above 2 mM initiate a response. For the remaining substrates (substrates 10 to 22 in Table 4), which exist exclusively within tissues, the net metabolic rate is

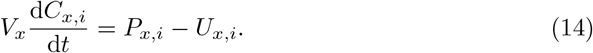

Production and utilization rates encompass all contributing metabolic reactions, formulated as:

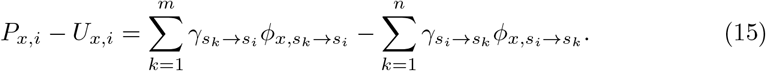

Here, 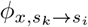 denotes the rate at which substrate *s*_*k*_ is utilized to form substrate *s*_*i*_ in tissue *x*, and vice versa for 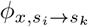 . The constants *γ*_(*·*)_ represent the respective stoichiometric coefficients. *m* refers to the number of processes forming substrate *i*, while *n* is the number of processes consuming substrate *i*.

### A.4 Basal metabolic reaction rate

Biochemical reactions *in vivo* are intricate metabolic processes that encompass multiple reaction steps. Our modeling approach follows a top-down systems perspective [29, 31, 33] and characterizes biochemical reactions as aggregate ‘pseudo’ processes by stoichiometrically combining several elementary reactions (Fig 15). Many lumped reactions are considered irreversible, reflecting the tendency of corresponding regulatory enzymes *in vivo* to have large equilibrium constants favoring product formation [33]. As part of our modelling framework, reversible reactions, such as the one involving the enzyme lactate dehydrogenase (LDH) where lactate is synthesized from pyruvate and can also be converted back to pyruvate, are deconstructed into two irreversible reactions. Consequently, all lumped reactions are treated as specific instances of a general irreversible, uni-uni (or bi-bi when two reactants are involved) substrate to product enzymatic reaction. Each reaction also includes the metabolic controller pairs ATP:ADP and NADH:NAD^+^, whose ratios are known to regulate reaction fluxes [33].

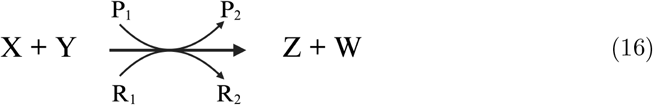

In this context, *P*_1_ and *P*_2_ denote ATP and ADP or vice versa, representing the phosphorylation state, while *R*_1_ and *R*_2_ stand for NADH and NAD^+^ or vice versa, indicating the redox state.

Rates of utilization 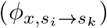 for any substrate *s*_*i*_ in tissue *x* vary based on the substrate concentration *C*_*x,i*_, unless otherwise specified, as well as on the phosphorylation state (PS) and the redox state (RS). As a general reaction, reactants *X* and *Y* and products *Z* and *W* are considered. The corresponding reaction rate equation in tissue *x* is:

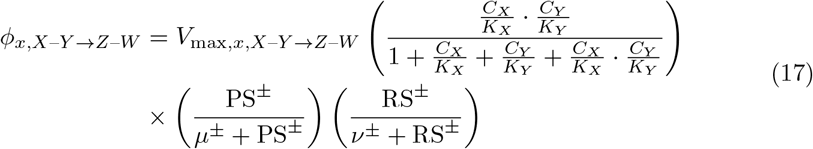

where *V*_max,*x,X*–*Y →Z*–*W*_, *K*_*X*_, and *K*_*Y*_ represent phenomenological maximum rate coefficients and Michaelis parameters specific to the reaction process. *C*_*X*_ and *C*_*Y*_ denote the concentrations of substrate *X* and *Y* in tissue *x*. The phosphorylation states are PS^+^ = *C*_ATP_*/C*_ADP_ in the forward direction and PS^*−*^ = *C*_ADP_*/C*_ATP_ in the opposite direction. Similarly, redox states are denoted as 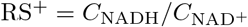 and 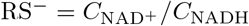. Parameters *μ*^*±*^ and *ν*^*±*^ are associated with the energy controller pairs.

It is noteworthy that AMP acts as a crucial allosteric regulator of glycogen phosphorylase for glycogenolysis (GLY*→*G6P) and phosphofructokinase-1 (PFK-1) for glycolysis II (G6P*→*GAP). Since AMP is not explicitly included in this model, AMP/ATP is approximated by [*ADP/ATP*]^2^ in these reactions (see Table S11, fluxes 2 and 8 in S1 Supporting Information). The maximum rate coefficients, Michaelis parameters, *μ*^*±*^ and *ν*^*±*^ values, along with in-tissue substrate concentrations, are sourced from Ref. [20] and the references therein.

### A.5 Metabolic response to eating

#### Postprandial metabolism and nutrient storage

Following a mixed meal, glucose concentrations in the systemic circulation typically increase within less than 30 minutes. However, it takes a longer interval, approximately 3-4 hours, to observe rises in blood fat levels [59]. The process of nutrient absorption is facilitated by a rapid increase in the insulin/glucagon ratio, which strongly influences postprandial metabolism [86, 87]. In healthy individuals, glucagon is suppressed after mixed meal or glucose ingestion, while insulin increases by at least an order of magnitude [54, 59, 88]. We thus simplified our approach by using insulin alone instead of the insulin-to-glucagon ratio as our metabolic signal from blood to tissues. Consequently, we define insulin-regulated enzyme activity factors, which are determined by plasma insulin concentration and vary for each organ:

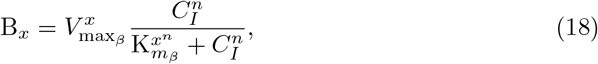

and

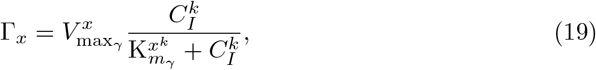

where activation by insulin is denoted by B_*x*_ and inhibition by insulin is denoted by Γ_*x*_. B_*x*_ and Γ_*x*_ are functions of insulin that are independent of substrate *i*. The subscript *x* denotes the different organs and tissues. The parameters 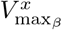 and 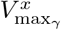 represent maximum rate coefficients, 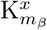 and 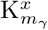 represent Michaelis parameters, *n* and *k* are hill coefficients modulating the strength of the response. For a given reaction, the maximum rate coefficient *V*_max,*x,i*_, characterizing the metabolic flux *i* in tissue *x* (see Eq 17), is modulated by insulin activity factors as follows:

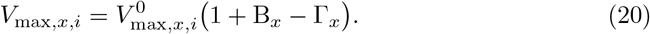

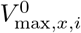 represents basal maximum rate coefficients (constants), while *V*_max,*x,i*_ represents dynamic maximum rate coefficients regulated by insulin, influencing metabolic rates via Eq (17). We note that for *V*_max,*x,i*_ to assume physiological values (*>* 0), 1 + B_*x*_ *>* Γ_*x*_ with B_*x*_, Γ_*x*_ *≥* 0. We also note that K_*m*_ values are chosen ‘large enough’ compared to (i.e., same order of magnitude as) postprandial insulin concentrations. As such, B_*x*_ ≈ 0 and Γ_*x*_ ≈ 0 when insulin returns to basal pre-meal levels. Table 6 lists the reactions modulated by insulin, along with the corresponding affected organs. Frayn [59] qualitatively described the enzymes and reactions affected by insulin. Consequently, we introduced insulin-regulated enzyme activity factors that modulate the production rates of the reactions outlined in Table 6.

**Table 6.** Reactions affected by pancreatic hormones during the postprandial phase.

| Hormone | Effect | Reaction # | Reaction | Affected organs |
| --- | --- | --- | --- | --- |
| Insulin | ↑ | 1 | Glycolysis I | H, M, G, L, A |
|  | ↑ | 2 | Glycolysis II | H, M, G, L, A |
|  | ↑ | 3 | Glycolysis III | H, M, G, L, A |
|  | ↑ | 7 | Glycogenesis | H, M, L |
|  | ↑ | 16 | Pyruvate oxidation | A, M, L |
|  | ↑ | 18 | Fatty acids synthesis | L |
|  | ↑ | 20 | TG synthesis | L, A |
|  | ↓ | 4 | Gluconeogenesis I | L |
|  | ↓ | 5 | Gluconeogenesis II | L |
|  | ↓ | 6 | Gluconeogenesis III | L |
|  | ↓ | 8 | Glycogenolysis | M, L |
|  | ↓ | 19 | Lipolysis | G, A |

#### Postabsorptive metabolism and nutrient mobilization

The term “postabsorptive state” implies that all contents of the previous meal have been absorbed from the GI tract, and that not much time has passed, as signs of starvation would otherwise appear. In the postabsorptive state, the blood glucose concentration usually hovers slightly below 5 mM. The concentration of insulin in plasma returns to basal values, which can vary widely among individuals but is typically around 60 pM. The concentration of glucagon also returns to basal values, approximately 20–30 pM [59]. Since insulin and glucagon concentrations are of comparable magnitudes, we employ the well-known glucagon/insulin ratio (GIR) and define the GIR-regulated enzyme activity factors that are active during the postabsorptive phase. This ratio correlates with metabolic changes during the postabsorptive phase [86].

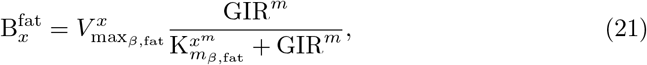

and

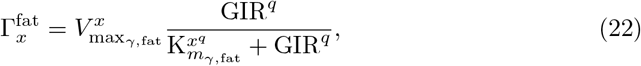

where activation by GIR is modulated by 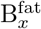 and inhibition by GIR is modulated by 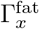. The subscript *x* denotes the different organs and tissues. The parameters 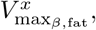 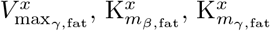, and exponents *m* and *q* are defined as before (see Eqs (18)-(19)). For a given reaction, the maximum rate coefficient *V*_max,*x,i*_ is modulated by GIR-regulated enzyme activity factors as follows:

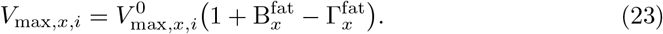

Here too, *V*_max,*x,i*_ assumes physiological values (*>* 0) when 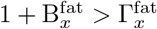 with 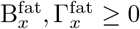. We note that 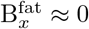 and 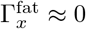 during the postprandial phase because GIR ≈ 0, i.e., *C*_*I*_ ≫ *C*_*G*_. Table 7 contains the reactions that are affected by insulin and glucagon, as well as which organ is affected.

**Table 7.** Reactions affected by pancreatic hormones during the postabsorptive phase.

| Hormone | Effect | Reaction # | Reaction | Affected organs |
| --- | --- | --- | --- | --- |
| Insulin | ↑ | 8 | Glycogenolysis | M |
|  | ↑ | 14 | Alanine formation | M |
|  | ↑ | 16 | Pyruvate oxidation | M |
|  | ↑ | 17 | Fatty acids oxidation | H, M |
|  | ↓ | 7 | Glycogenesis | M |
| Glucagon | ↑ | 4 | Gluconeogenesis I | L |
|  | ↑ | 5 | Gluconeogenesis II | L |
|  | ↑ | 6 | Gluconeogenesis III | L |
|  | ↑ | 8 | Glycogenolysis | L |
|  | ↑ | 10 | Lactate oxidation | L |
|  | ↑ | 15 | Alanine utilization | L |
|  | ↑ | 16 | Pyruvate oxidation | L |
|  | ↑ | 17 | Fatty acids oxidation | L |
|  | ↓ | 7 | Glycogenesis | L |
|  | ↓ | 9 | Pyruvate reduction | L |

Notably, skeletal muscle and cardiac cells lack glucagon receptors [89], but reduced insulin levels can signal metabolic pathways in these organs during the postabsorptive phase [90]. Thus, during the postabsorptive phase, the heart and skeletal muscle solely respond to an insulin signal:

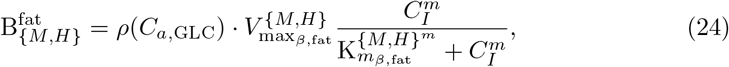

and

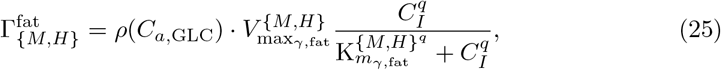

where

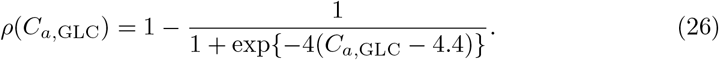

The logistic sigmoid function, denoted as *ρ*(*C*_*a*,GLC_), serves as a switch governing the activation and deactivation of postabsorptive insulin-regulated enzyme activity factors in skeletal muscle and heart. It is based on blood glucose levels. The threshold defining the shift from normoglycemia to hypoglycemia is established at 4.4 mM, as specified by Berger and Zdzieblo [78]. This value is utilized to determine the initiation of the postabsorptive phase in these organs. It is noteworthy that *ρ*(*C*_*a*,GLC_) approaches 0 during the postprandial phase when *C*_*a*,GLC_ *>* 4.4 mM, and so is only activated during the postabsorptive phase. The slope, set to *−*4, ensures a rapid shift from activation (*ρ*(*C*_*a*,GLC_) ≈ 1) to deactivation (*ρ*(*C*_*a*,GLC_) ≈ 0) even with slight deviations of *C*_*a*,GLC_ above 4.4 mM. This modeling decision is influenced by the observation that skeletal myocytes can directly sense extracellular glucose concentrations, activating a signaling pathway for hormone-independent metabolic activation. This glucose sensing pathway acts in concert with insulin to enhance muscle glucose utilization, contributing to the maintenance of systemic glucose homeostasis [86, 91]. Table 7 lists the reactions in skeletal muscle and heart that are affected by insulin during the postabsorptive phase.

### A.6 Virtual subjects

We consider lean, healthy male and female subjects. Table 8 provides details regarding physical attributes, including tissue weights and basal blood flows.

**Table 8.**
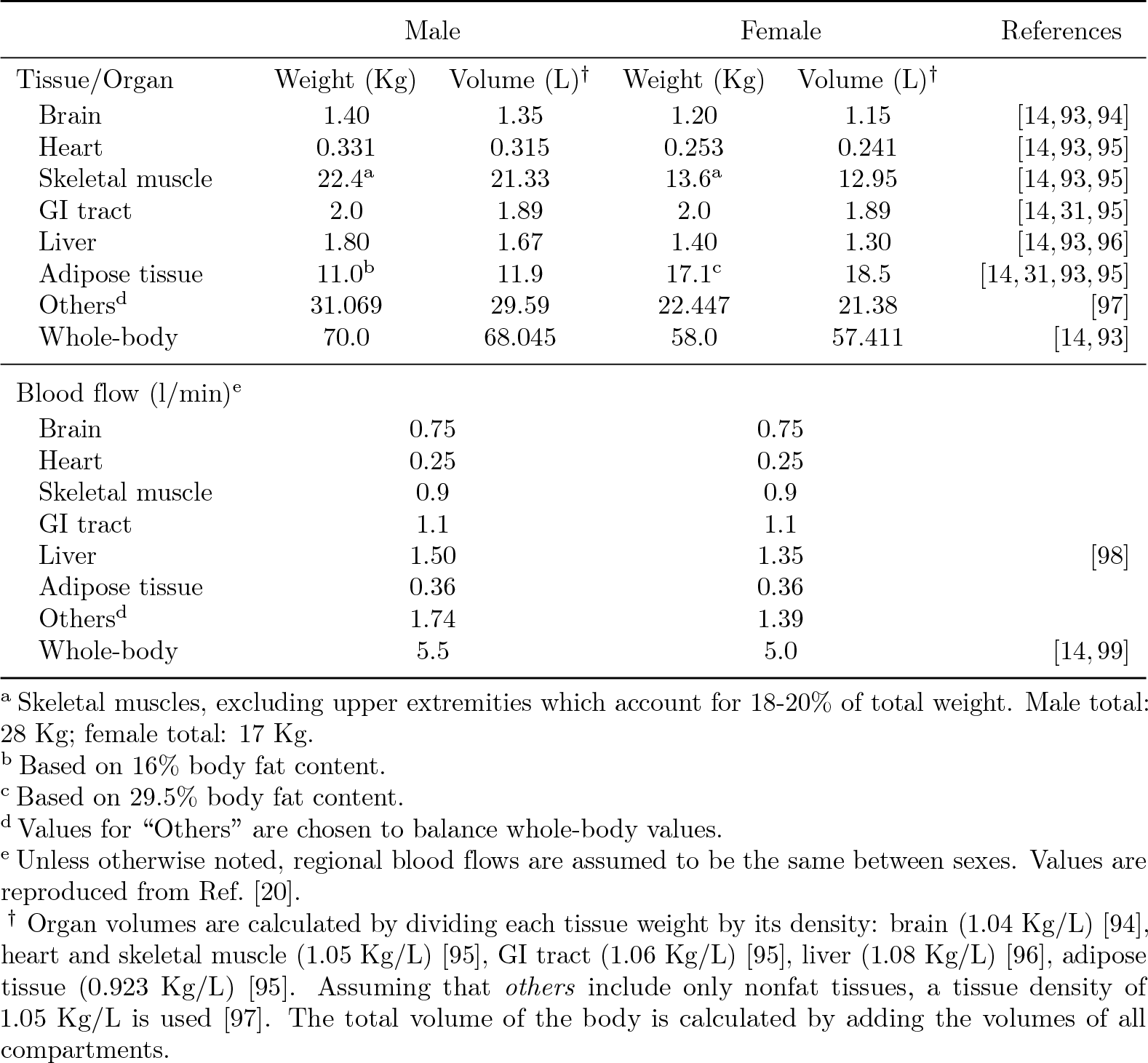
Physical characteristics.

|  | Male |  | Female |  | References |
| --- | --- | --- | --- | --- | --- |
| Tissue/Organ | Weight (Kg) | Volume (L)<sup>†</sup> | Weight (Kg) | Volume (L)<sup>†</sup> |  |
| Brain | 1.40 | 1.35 | 1.20 | 1.15 | [14, 93, 94] |
| Heart | 0.331 | 0.315 | 0.253 | 0.241 | [14, 93, 95] |
| Skeletal muscle | 22.4<sup>a</sup> | 21.33 | 13.6<sup>a</sup> | 12.95 | [14, 93, 95] |
| GI tract | 2.0 | 1.89 | 2.0 | 1.89 | [14, 31, 95] |
| Liver | 1.80 | 1.67 | 1.40 | 1.30 | [14, 93, 96] |
| Adipose tissue | 11.0<sup>b</sup> | 11.9 | 17.1<sup>c</sup> | 18.5 | [14, 31, 93, 95] |
| Others<sup>d</sup> | 31.069 | 29.59 | 22.447 | 21.38 | [97] |
| Whole-body | 70.0 | 68.045 | 58.0 | 57.411 | [14, 93] |
| Blood flow (l/min)<sup>e</sup> |  |  |  |  |  |
| Brain | 0.75 |  | 0.75 |  |  |
| Heart | 0.25 |  | 0.25 |  |  |
| Skeletal muscle | 0.9 |  | 0.9 |  |  |
| GI tract | 1.1 |  | 1.1 |  |  |
| Liver | 1.50 |  | 1.35 |  | [98] |
| Adipose tissue | 0.36 |  | 0.36 |  |  |
| Others<sup>d</sup> | 1.74 |  | 1.39 |  |  |
| Whole-body | 5.5 |  | 5.0 |  | [14, 99] |

<sup>a</sup> Skeletal muscles, excluding upper extremities which account for 18-20% of total weight. Male total: 28 Kg; female total: 17 Kg.
<sup>b</sup> Based on 16% body fat content.
<sup>c</sup> Based on 29.5% body fat content.
<sup>d</sup> Values for “Others” are chosen to balance whole-body values.
<sup>e</sup> Unless otherwise noted, regional blood flows are assumed to be the same between sexes. Values are reproduced from Ref. [20].
<sup>†</sup> Organ volumes are calculated by dividing each tissue weight by its density: brain (1.04 Kg/L) [94], heart and skeletal muscle (1.05 Kg/L) [95], GI tract (1.06 Kg/L) [95], liver (1.08 Kg/L) [96], adipose tissue (0.923 Kg/L) [95]. Assuming that *others* include only nonfat tissues, a tissue density of 1.05 Kg/L is used [97]. The total volume of the body is calculated by adding the volumes of all compartments.

<u>Male:</u> 70 Kg body weight, aged between 20 and 35 years. Regarding body composition, skeletal muscle constitutes 40% of the total body weight, and adipose tissue accounts for 16%. In the initial conditions for each simulation, the subject is in a state of overnight fast (8-12 h), with a cardiac output of 5.5 l/min and a resting RQ of 0.8.

<u>Female:</u> 58 Kg body weight, aged between 20 and 35 years. Regarding body composition, skeletal muscle constitutes 30% of the total body weight, and adipose tissue accounts for 29.5%. In the initial conditions for each simulation, the subject is in a state of overnight fast (8-12 h), with a cardiac output of 5 l/min and a resting RQ of 0.8.

## Appendix B Parameter estimation

While the qualitative effects of insulin and glucagon are known (Tables 6-7), the specific parameters governing insulin- and glucagon-induced metabolic regulation remain less established. We conducted parameter estimations based on experimental data [54–56, 58] and reference ranges from literature sources (Table 9). The parameter values for simulating the metabolism of a healthy adult (either male or female) in an overnight fasted condition are detailed in our prior work [20] and are also accessible in the S1 Supporting Information. Parameters associated with metabolic changes in response to feeding and subsequent fasting, as outlined in Table 1, were estimated using *fmincon*, a gradient-based constrained optimization algorithm commonly known as *constrained nonlinear optimization* or *nonlinear programming*. This algorithm was applied to minimize the difference between model-predicted concentrations and concentration data over 6 to 10 hours following various mixed meals.

**Table 9.**
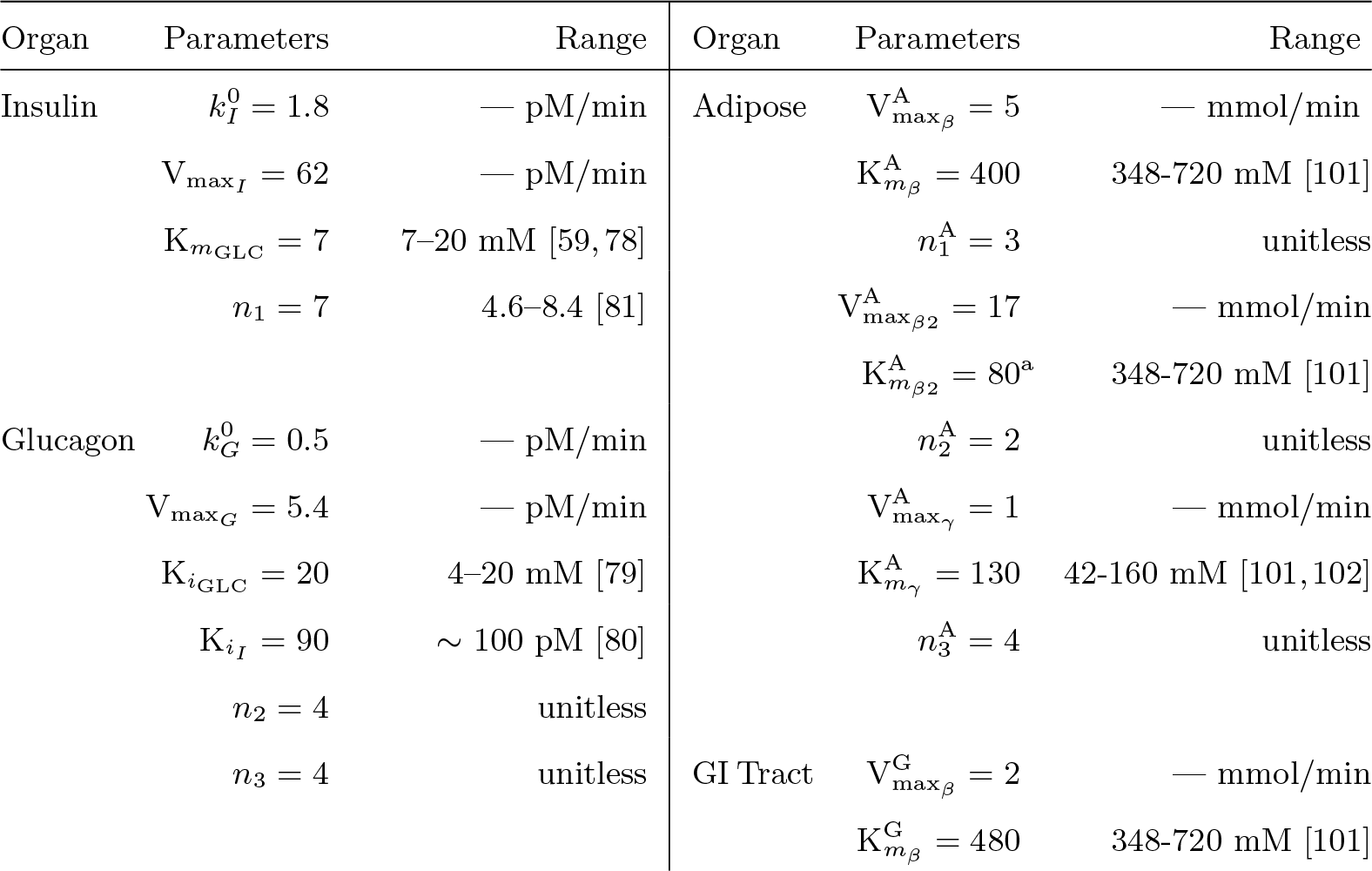
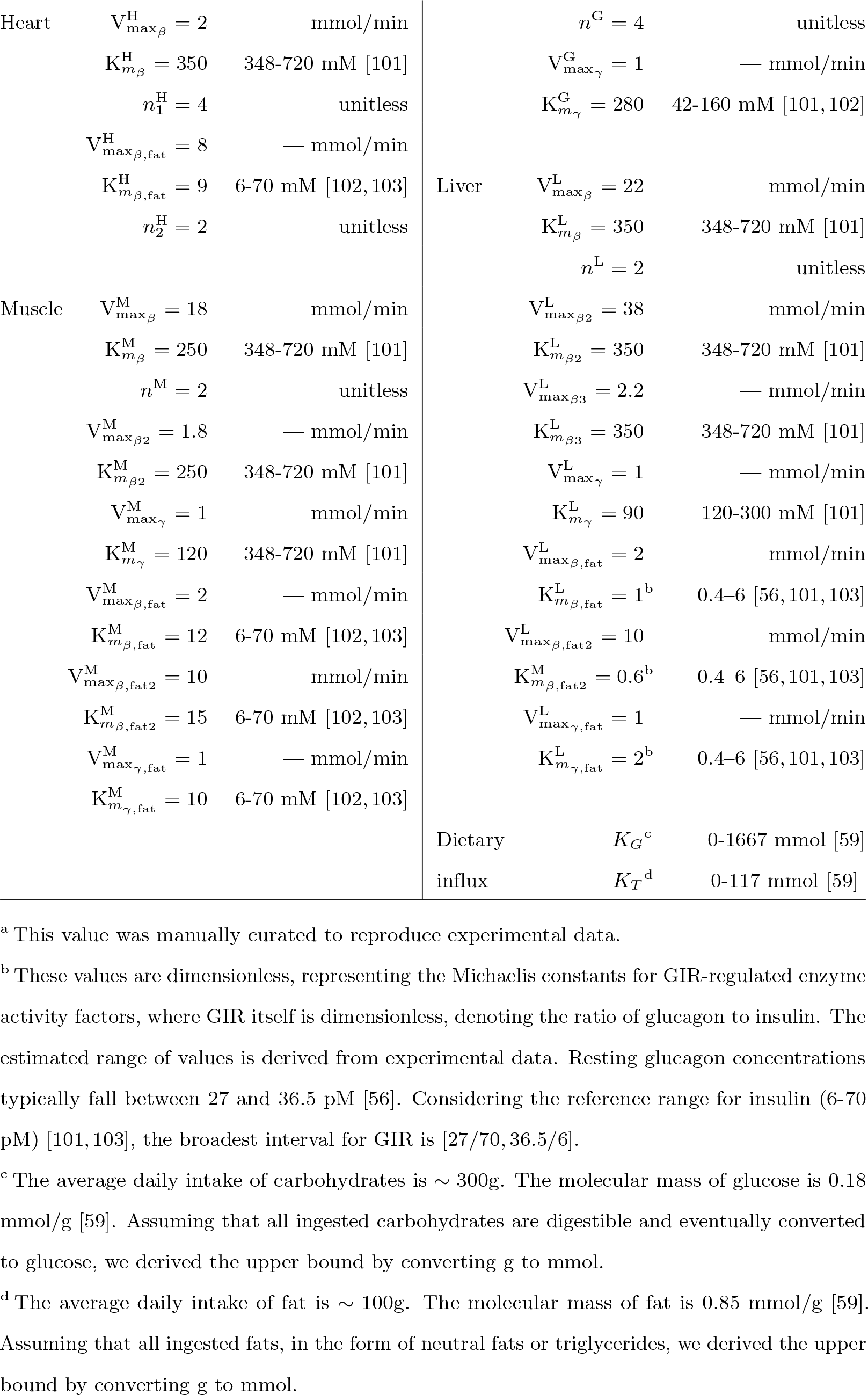
Estimated model parameters.

Let *p*_0_ denote the initial guess set for model parameters. For parameters associated with in-tissue regulation by insulin and glucagon (18–25), the maximum rate coefficients 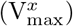 for enzyme activity factors are fixed at 1. The Michaelis constants 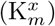 are set to the midpoint of the reference range obtained from the literature, while the hill coefficients (*n*^*x*^) are set to 1. Concerning the dynamics of plasma insulin and glucagon (7–8), the maximum rate coefficients 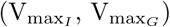, basal secretion rates 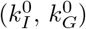, and hill coefficients (*n*_2_, *n*_3_) are uniformly set to 1. Regarding dietary influx (5–6), the half-emptying gut thresholds for glucose (*K*_*G*_) and TG (*K*_*T*_) are fixed at half the values of the average daily intake of macronutrients, as outlined in the literature. Reference ranges are elaborated in Table 9.

We then established a set of lower (lb) and upper (ub) bounds for the free variables in *p*_0_, ensuring that the solution remains within the range lb ≤ *p*_0_ ≤ ub. Hill coefficients are confined to the range of 1 to 4, unless specific evidence suggests otherwise, given that cooperative binding and allosteric enzymatic processes seldom produce Hill coefficients exceeding 4 [100]. All half-maximum constants (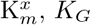, and *K*_*T*_) have lb = min(range)*/*2 and ub = max(range) *×* 2. We opted not to strictly adhere to experimental ranges to accommodate variability between experiments and acknowledge that many reactions are modeled as ‘pseudo’ processes through stoichiometric combinations of several elementary reactions (Fig 15). Maximum rate coefficients 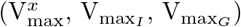 are restricted between 0 and 100 when modeling the activation of a reaction and between 0 and 1 when modeling the inhibition of a reaction. The latter constraint arises from the equation form for inhibition (see Eqs 20 and 23). Model equations are solved using *ode15s* (MATLAB 2021a), a variable-step, variable-order solver ranging from orders 1 to 5. This solver is an implicit integration algorithm designed for stiff systems.

For a given substrate (or hormone) *s*, consider the weighted residuals:

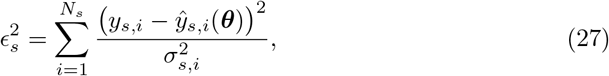

where *y*_*s,i*_ represents the measured data point *i* for substrate (or hormone) *s, σ*_*s,i*_ denotes the corresponding experimental standard error, *N*_*s*_ signifies the total number of data points for substrate *s*, and ŷ _*s,i*_(***θ***) is the predicted value given the set of parameters ***θ***. Consequently, we aim to minimize a problem specified by

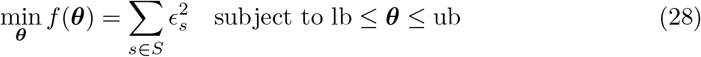

Set *S* = *{*insulin, glucagon, GLC, LAC, TG, FFA, GLR, GLY_M_, GLY_L_*}* specifically for the male model. The subscripts M and L correspond to skeletal muscle and liver, respectively. Substrate abbreviations are outlined in Table 4. The data represent plasma concentrations of hormones and substrates and are detailed in Table 10. We utilized time series data reflecting dynamic concentrations following single meals of various compositions spanning 6 to 10 hours post food intake. All datasets were concurrently employed for parameter estimation. A comprehensive list of estimated parameters is provided in Table 9.

**Table 10.**
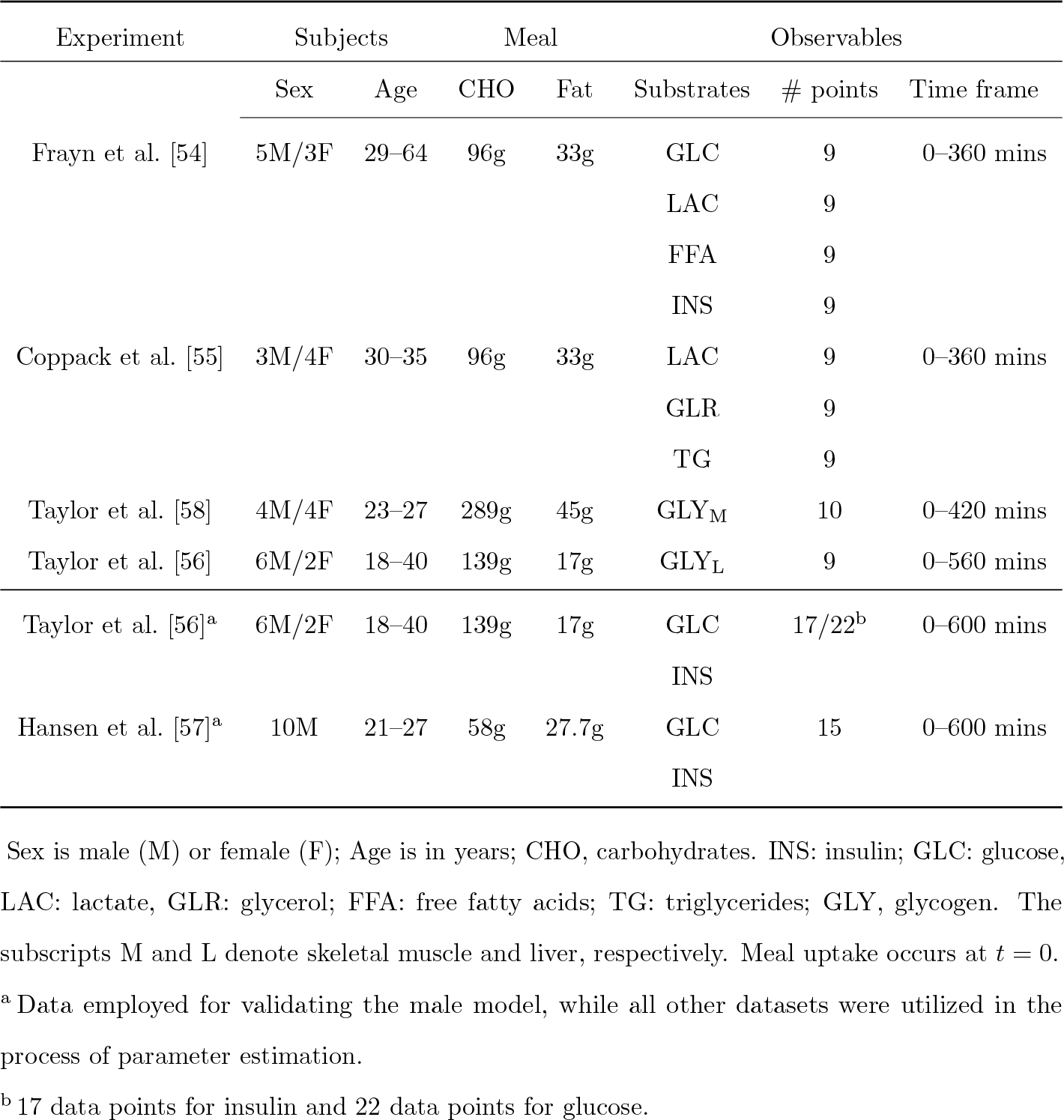
Experimental data used for parameter estimation and model validation.

## Supporting information

**S1 Supporting Information Model equations, parameters, and sensitivity analysis results**. The file contains the following sections: Section 1: Model parameters; Section 2: Sensitivity analysis; Section 3: Model equations; Section 4: Key metabolic fluxes; Section 5: Code availability.

**Fig 1 in S1 Supporting Information. Results of local sensitivity analysis at 3 hours post-meal**. (a) Male model and HiC; (b) female model and HiC; (c) male model and HiF; (d) Female model and HiF. HiC, high-carbohydrate meal; HiF, high-fat meal. Glycolysis II, *ϕ*_G6P*→*GAP_; gluconeogenesis II, *ϕ*_GAP*→*G6P_; glycogenesis, *ϕ*_G6P*→*GLY_; glycogenolysis, *ϕ*_GLY*→*G6P_, lipolysis, *ϕ*_TG*→*FFA–GLR_.

**Fig 2 in S1 Supporting Information. Results of local sensitivity analysis at 9 hours post-meal**. (a) Male model and HiC; (b) female model and HiC; (c) male model and HiF; (d) Female model and HiF. HiC, high-carbohydrate meal; HiF, high-fat meal. Glycolysis II, *ϕ*_G6P*→*GAP_; gluconeogenesis II, *ϕ*_GAP*→*G6P_; glycogenesis, *ϕ*_G6P*→*GLY_; glycogenolysis, *ϕ*_GLY*→*G6P_, lipolysis, *ϕ*_TG*→*FFA–GLR_.

**Fig 3 in S1 Supporting Information. Results of local sensitivity analysis at 24 hours post-meal**. (a) Male model and HiC; (b) female model and HiC; (c) male model and HiF; (d) Female model and HiF. HiC, high-carbohydrate meal; HiF, high-fat meal. Glycolysis II, *ϕ*_G6P*→*GAP_; gluconeogenesis II, *ϕ*_GAP*→*G6P_; glycogenesis, *ϕ*_G6P*→*GLY_; glycogenolysis, *ϕ*_GLY*→*G6P_, lipolysis, *ϕ*_TG*→*FFA–GLR_.

## Acknowledgments

Anita T. Layton is supported by the Canada 150 Research Chairs Program, the Natural Sciences Canadian Institutes of Health Research (CIHR), and the Natural Sciences and Engineering Research Council of Canada (NSERC) Discovery award (RGPIN-2019-03916).

## References

1. NCD Risk Factor Collaboration (NCD-RisC). Trends in adult body-mass index in 200 countries from 1975 to 2014: a pooled analysis of 1698 population-based measurement studies with 19·2 million participants. Lancet. 2016;10026(387):1377–1396. doi:10.1016/S0140-6736(16)30054-X.

2. Cho NH, Shaw J, Karuranga S, Huang Y, da Rocha Fernandes J, Ohlrogge A, et al. IDF Diabetes Atlas: Global estimates of diabetes prevalence for 2017 and projections for 2045. Diabetes Res Clin Pract. 2018;138:271–281. doi:10.1016/j.diabres.2018.02.023.

3. Devries MC, Lowther SA, Glover AW, Hamadeh MJ, Tarnopolsky MA. IMCL area density, but not IMCL utilization, is higher in women during moderate-intensity endurance exercise, compared with men. Am J Physiol Regul Integr Comp Physiol. 2007;293(6):R2336–R2342. doi:10.1152/ajpregu.00510.2007.

4. Beaudry KM, Devries MC. Sex-based differences in hepatic and skeletal muscle triglyceride storage and metabolism. Appl Physiol Nutr Metab. 2019;44(8):805–813. doi:10.1139/apnm-2018-0635.

5. Bickerton A, Roberts R, Fielding B, Tornqvist H, Blaak E, Wagenmakers A, et al. Adipose tissue fatty acid metabolism in insulin-resistant men. Diabetologia. 2008;51:1466–1474. doi:10.1007/s00125-008-1040-x.

6. Hedrington MS, Davis SN. Sexual dimorphism in glucose and lipid metabolism during fasting, hypoglycemia, and exercise. Front Endocrinol. 2015;6:61. doi:10.3389/fendo.2015.00061.

7. Varlamov O, Bethea CL, Roberts Jr CT. Sex-specific differences in lipid and glucose metabolism. Front Endocrinol. 2015;5:241. doi:10.3389/fendo.2014.00241.

8. Tarnopolsky MA, Rennie CD, Robertshaw HA, Fedak-Tarnopolsky SN, Devries MC, Hamadeh MJ. Influence of endurance exercise training and sex on intramyocellular lipid and mitochondrial ultrastructure, substrate use, and mitochondrial enzyme activity. Am J Physiol Regul Integr Comp Physiol. 2007;292(3):R1271–R1278. doi:10.1152/ajpregu.00472.2006.

9. Oraha J, Enriquez RF, Herzog H, Lee NJ. Sex-specific changes in metabolism during the transition from chow to high-fat diet feeding are abolished in response to dieting in C57BL/6J mice. Int J Obes. 2022;46(10):1749–1758. doi:10.1038/s41366-022-01174-4.

10. Santosa S, Jensen MD. The sexual dimorphism of lipid kinetics in humans. Front Endocrinol. 2015;6:103. doi:10.3389/fendo.2015.00103.

11. Corssmit E, Stouthard J, Romijn J, Endert E, Sauerwein H. Sex differences in the adaptation of glucose metabolism to short-term fasting: effects of oral contraceptives. Metabolism. 1994;43(12):1503–1508. doi:10.1016/0026-0495(94)90008-6.

12. Pramfalk C, Pavlides M, Banerjee R, McNeil CA, Neubauer S, Karpe F, et al. Sex-specific differences in hepatic fat oxidation and synthesis may explain the higher propensity for NAFLD in men. J Clin Endocrinol Metab. 2015;100(12):4425–4433. doi:10.1210/jc.2015-2649.

13. Lebeck J. Sexual dimorphism in Glucose and lipid Metabolism; 2016. Available from: 10.3389/fendo.2016.00166.

14. Thiele I, Sahoo S, Heinken A, Hertel J, Heirendt L, Aurich MK, et al. Personalized whole-body models integrate metabolism, physiology, and the gut microbiome. Mol Syst Biol. 2020;16(5):e8982. doi:10.15252/msb.20198982.

15. Leone A, De Amicis R, Bertoli S, Spadafranca A, De Carlo G, Battezzati A. Absence of a sexual dimorphism in postprandial glucose metabolism after administration of a balanced mixed meal in healthy young volunteers. Nutr Diabetes. 2022;12(1):6. doi:10.1038/s41387-022-00184-5.

16. Carter S, Rennie C, Tarnopolsky M. Substrate utilization during endurance exercise in men and women after endurance training. Am J Physiol Endocrinol Metab. 2001;doi:10.1152/ajpendo.2001.280.6.E898.

17. Mittendorfer B, Horowitz JF, Klein S. Gender differences in lipid and glucose kinetics during short-term fasting. Am J Physiol Endocrinol Metab. 2001;281(6):E1333–E1339. doi:10.1152/ajpendo.2001.281.6.E1333.

18. Davis SN, Galassetti P, Wasserman DH, Tate D. Effects of gender on neuroendocrine and metabolic counterregulatory responses to exercise in normal man. J Clin Endocrinol Metab. 2000;85(1):224–230. doi:10.1210/jcem.85.1.6328.

19. Sandoval DA, D’Alessio DA. Physiology of proglucagon peptides: role of glucagon and GLP-1 in health and disease. Physiol Rev. 2015;95(2):513–548. doi:10.1152/physrev.00013.2014.

20. Abo SM, Casella E, Layton AT. Sexual Dimorphism in Substrate Metabolism During Exercise. Bull Math Biol. 2024;86(2):17. doi:10.1007/s11538-023-01242-4.

21. Palumbo MC, Morettini M, Tieri P, Diele F, Sacchetti M, Castiglione F. Personalizing physical exercise in a computational model of fuel homeostasis. PLoS Comput Biol. 2018;14(4):e1006073. doi:10.1371/journal.pcbi.1006073.

22. Qiu S, Vazquez JT, Boulger E, Liu H, Xue P, Hussain MA, et al. Hepatic estrogen receptor α is critical for regulation of gluconeogenesis and lipid metabolism in males. Sci Rep. 2017;7(1):1661. doi:10.1038/s41598-017-01937-4.

23. Palmisano BT, Zhu L, Stafford JM. In: Mauvais-Jarvis F, editor. Role of Estrogens in the Regulation of Liver Lipid Metabolism. Cham: Springer International Publishing; 2017. p. 227–256.

24. Shen M, Shi H, et al. Sex hormones and their receptors regulate liver energy homeostasis. Int J Endocrinol. 2015;2015. doi:10.1155/2015/294278.

25. Hunter SK. Sex differences in human fatigability: mechanisms and insight to physiological responses. Acta Physiol (oxf). 2014;210(4):768–789. doi:10.1111/apha.12234.

26. Tarnopolsky L, MacDougall J, Atkinson S, Tarnopolsky M, Sutton J. Gender differences in substrate for endurance exercise. J Appl Physiol. 1990;68(1):302–308. doi:10.1152/jappl.1990.68.1.302.

27. Tarnopolsky M, Atkinson S, Phillips S, MacDougall J. Carbohydrate loading and metabolism during exercise in men and women. J Appl Physiol. 1995;78(4):1360–1368. doi:10.1152/jappl.1995.78.4.1360.

28. Cheneviere X, Borrani F, Sangsue D, Gojanovic B, Malatesta D. Gender differences in whole-body fat oxidation kinetics during exercise. Appl Physiol Nutr Metab. 2011;36(1):88–95. doi:10.1139/H10-086.

29. Kurata H. Virtual metabolic human dynamic model for pathological analysis and therapy design for diabetes. iScience. 2021;24(2). doi:10.1016/j.isci.2021.102101.

30. Carstensen PE, Bendsen J, Reenberg AT, Ritschel TK, Jørgensen JB. A whole-body multi-scale mathematical model for dynamic simulation of the metabolism in man. IFAC Pap OnLine. 2022;55(23):58–63. doi:10.1016/j.ifacol.2023.01.015.

31. Kim J, Saidel GM, Cabrera ME. Multi-Scale Computational Model of Fuel Homeostasis During Exercise: Effect of Hormonal Control. Ann Biomed Eng. 2006;doi:10.1007/s10439-006-9201-x.

32. König M, Bulik S, Holzhütter HG. Quantifying the Contribution of the Liver to Glucose Homeostasis: A Detailed Kinetic Model of Human Hepatic Glucose Metabolism. PLOS Comput Biol. 2012;8(6):1–17. doi:10.1371/journal.pcbi.1002577.

33. Dash RK, DiBella JA, Cabrera ME. A computational model of skeletal muscle metabolism linking cellular adaptations induced by altered loading states to metabolic responses during exercise. Biomed Eng Online. 2007;6(1):1–28. doi:10.1186/1475-925X-6-14.

34. Berndt N, Holzhütter HG. Dynamic metabolic zonation of the hepatic glucose metabolism is accomplished by sinusoidal plasma gradients of nutrients and hormones. Front Physiol. 2018;9:1786. doi:10.3389/fphys.2018.01786.

35. Berndt N, Bulik S, Wallach I, Wünsch T, König M, Stockmann M, et al. HEPATOKIN1 is a biochemistry-based model of liver metabolism for applications in medicine and pharmacology. Nat Commun. 2018;9(1):2386. doi:10.1038/s41467-018-04720-9.

36. Li Y, Solomon TP, Haus JM, Saidel GM, Cabrera ME, Kirwan JP. Computational model of cellular metabolic dynamics: effect of insulin on glucose disposal in human skeletal muscle. Am J Physiol Endocrinol Metab. 2010;298(6):E1198–E1209. doi:10.1152/ajpendo.00713.2009.

37. Sluka JP, Fu X, Swat M, Belmonte JM, Cosmanescu A, Clendenon SG, et al. A liver-centric multiscale modeling framework for xenobiotics. PLoS One. 2016;11(9):e0162428. doi:10.1371/journal.pone.0162428.

38. Ashworth WB, Davies NA, Bogle IDL. A computational model of hepatic energy metabolism: understanding zonated damage and steatosis in NAFLD. PLoS Comput Biol. 2016;12(9):e1005105. doi:10.1371/journal.pcbi.1005105.

39. Cobelli C, Renard E, Kovatchev B. Artificial pancreas: past, present, future. Diabetes. 2011;60(11):2672–2682. doi:10.2337/db11-0654.

40. Visentin R, Campos-Náñez E, Schiavon M, Lv D, Vettoretti M, Breton M, et al. The UVA/Padova type 1 diabetes simulator goes from single meal to single day. J Diabetes Sci Technol. 2018;12(2):273–281. doi:10.1177/1932296818757747.

41. Pearson T, Wattis JA, King J, MacDonald IA, Mazzatti D. A mathematical model of the human metabolic system and metabolic flexibility. Bull Math Biol. 2014;76:2091–2121. doi:10.1007/s11538-014-0001-4.

42. Hetherington J, Sumner T, Seymour R, Li L, Rey MV, Yamaji S, et al. A composite computational model of liver glucose homeostasis. I. Building the composite model. J R Soc Interf. 2012;9(69):689–700. doi:10.1098/rsif.2011.0141.

43. Pratt AC, Wattis JA, Salter AM. Mathematical modelling of hepatic lipid metabolism. Math Biosci. 2015;262:167–181. doi:10.1016/j.mbs.2014.12.012.

44. Sweatman CZH. Mathematical model of diabetes and lipid metabolism linked to diet, leptin sensitivity, insulin sensitivity and VLDLTG clearance predicts paths to health and type II diabetes. J Theor Biol. 2020;486:110037. doi:10.1016/j.jtbi.2019.110037.

45. Panunzi S, Pompa M, Borri A, Piemonte V, De Gaetano A. A revised Sorensen model: Simulating glycemic and insulinemic response to oral and intra-venous glucose load. PLoS One. 2020;15(8):e0237215. doi:10.1371/journal.pone.0237215.

46. Sorensen JT. A physiologic model of glucose metabolism in man and its use to design and assess improved insulin therapies for diabetes; 1985.

47. Yasemi M, Jolicoeur M. Modelling cell metabolism: a review on constraint-based steady-state and kinetic approaches. Processes. 2021;9(2):322. doi:10.3390/pr9020322.

48. Cvitanović Tomaš T, Urlep Ž, Moškon M, Mraz M, Rozman D. LiverSex computational model: sexual aspects in hepatic metabolism and abnormalities. Front Physiol. 2018;9:360. doi:10.3389/fphys.2018.00360.

49. Swapnasrita S, Carlier A, Layton AT. Sex-specific computational models of kidney function in patients with diabetes. Front Physiol. 2022;13:14. doi:10.3389/fphys.2022.741121.

50. Sekizkardes H, Chung ST, Chacko S, Haymond MW, Startzell M, Walter M, et al. Free fatty acid processing diverges in human pathologic insulin resistance conditions. J Clin Invest. 2020;130(7):3592–3602. doi:10.1172/JCI135431.

51. Staehr P, Hother-Nielsen O, Landau BR, Chandramouli V, Holst JJ, Beck-Nielsen H. Effects of free fatty acids per se on glucose production, gluconeogenesis, and glycogenolysis. Diabetes. 2003;52(2):260–267. doi:10.2337/diabetes.52.2.260.

52. Chen X, Iqbal N, Boden G, et al. The effects of free fatty acids on gluconeogenesis and glycogenolysis in normal subjects. J Clin Invest. 1999;103(3):365–372. doi:10.1172/JCI5479.

53. Browning JD, Baxter J, Satapati S, Burgess SC. The effect of short-term fasting on liver and skeletal muscle lipid, glucose, and energy metabolism in healthy women and men. J Lipid Res. 2012;53(3):577–586. doi:10.1194/jlr.P020867.

54. Frayn KN, Coppack SW, Humphreys SM, Clark ML, Evans RD. Periprandial regulation of lipid metabolism in insulin-treated diabetes mellitus. Metabolism. 1993;42(4):504–510. doi:10.1016/0026-0495(93)90110-A.

55. Coppack S, Fisher R, Gibbons G, Humphreys S, McDonough M, Potts J, et al. Postprandial substrate deposition in human forearm and adipose tissues in vivo. Clin Sci. 1990;79(4):339–348. doi:10.1042/cs0790339.

56. Taylor R, Magnusson I, Rothman DL, Cline GW, Caumo A, Cobelli C, et al. Direct assessment of liver glycogen storage by 13C nuclear magnetic resonance spectroscopy and regulation of glucose homeostasis after a mixed meal in normal subjects. J Clin Invest. 1996;97(1):126–132. doi:10.1172/JCI118379.

57. Hansen KB, Vilsbøll T, Bagger JI, Holst JJ, Knop FK. Increased postprandial GIP and glucagon responses, but unaltered GLP-1 response after intervention with steroid hormone, relative physical inactivity, and high-calorie diet in healthy subjects. J Clin Endocrinol Metab. 2011;96(2):447–453. doi:10.1210/jc.2010-160.

58. Taylor R, Price TB, Katz L, Shulman RG, Shulman GI. Direct measurement of change in muscle glycogen concentration after a mixed meal in normal subjects. Am J Physiol Endocrinol Metab. 1993;265(2):E224–E229. doi:10.1152/ajpendo.1993.265.2.E224.

59. Frayn KN. 2–7. In: Metabolic regulation: A Human Perspective. 3rd ed. Oxford, United Kingdom: John Wiley & Sons; 2010. p. 27–168.

60. Ludwig DS. The glycemic index: physiological mechanisms relating to obesity, diabetes, and cardiovascular disease. Jama. 2002;287(18):2414–2423. doi:10.1001/jama.287.18.2414.

61. Hiyoshi T, Fujiwara M, Yao Z. Postprandial hyperglycemia and postprandial hypertriglyceridemia in type 2 diabetes. J Biomed Res. 2019;33(1):1. doi:10.7555/JBR.31.20160164.

62. Lundsgaard AM, Kiens B. Gender differences in skeletal muscle substrate metabolism–molecular mechanisms and insulin sensitivity. Front Endocrinol. 2014;5:195. doi:10.3389/fendo.2014.00195.

63. Ivey PA, Gaesser GA. Postexercise muscle and liver glycogen metabolism in male and female rats. J Appl Physiol. 1987;62(3):1250–1254. doi:10.1152/jappl.1987.62.3.1250.

64. Jensen J, Rustad PI, Kolnes AJ, Lai YC. The role of skeletal muscle glycogen breakdown for regulation of insulin sensitivity by exercise. Front Physiol. 2011;2:112. doi:10.3389/fphys.2011.00112.

65. Emhoff CAW, Messonnier LA. Concepts of lactate metabolic clearance rate and lactate clamp for metabolic inquiry: a mini-review. Nutrients. 2023;15(14):3213. doi:10.3390/nu15143213.

66. Hugi D, Bracher V, Tappy L, Blum J. Postprandial hydrogen breath excretion, plasma lactate concentration, glucose metabolism and insulin levels in veal calves. J Anim Physiol Anim Nutr. 1997;78(1-5):42–48. doi:10.1111/j.1439-0396.1997.tb00854.x.

67. Jackson R, Hamling J, Sim B, Hawa M, Blix P, Nabarro J. Peripheral lactate and oxygen metabolism in man: the influence of oral glucose loading. Metabolism. 1987;36(2):144–150. doi:10.1016/0026-0495(87)90008-4.

68. Carroll MD, Lacher DA, Sorlie PD, Cleeman JI, Gordon DJ, Wolz M, et al. Trends in serum lipids and lipoproteins of adults, 1960-2002. JAMA. 2005;294(14):1773–1781. doi:10.1001/jama.294.14.1773.

69. Ramos-Jiménez A, Hernández-Torres RP, Torres-Durán PV, Romero-Gonzalez J, Mascher D, Posadas-Romero C, et al. The respiratory exchange ratio is associated with fitness indicators both in trained and untrained men: a possible application for people with reduced exercise tolerance. Clin Med Circ Respirat Pulm Med. 2008;2:CCRPM–S449. doi:10.4137/ccrpm.s449.

70. Deuster PA, Heled Y. Chapter 41 - Testing for Maximal Aerobic Power. In: Seidenberg PH, Beutler AI, editors. The Sports Medicine Resource Manual. Philadelphia: W.B. Saunders; 2008. p. 520–528. Available from: 10.1016/B978-141603197-0.10069-2.

71. Roepstorff C, Steffensen CH, Madsen M, Stallknecht B, Kanstrup IL, Richter EA, et al. Gender differences in substrate utilization during submaximal exercise in endurance-trained subjects. Am J Physiol Endocrinol Metab. 2002;282(2):E435–E447. doi:10.1152/ajpendo.00266.2001.

72. Whytock KL, Shepherd SO, Cocks M, Wagenmakers AJ, Strauss JA. Young, healthy males and females present cardiometabolic protection against the detrimental effects of a 7-day high-fat high-calorie diet. Eur J Nutr. 2021;60:1605–1617. doi:10.1007/s00394-020-02357-3.

73. Brooks GA, Curl CC, Leija RG, Osmond AD, Duong JJ, Arevalo JA. Tracing the lactate shuttle to the mitochondrial reticulum. Exp Mol Med. 2022;54(9):1332–1347. doi:10.1038/s12276-022-00802-3.

74. Legouis D, Faivre A, Cippà PE, de Seigneux S. Renal gluconeogenesis: an underestimated role of the kidney in systemic glucose metabolism. Nephrol Dial Transplant. 2022;37(8):1417–1425. doi:10.1093/ndt/gfaa302.

75. Puskarich MA, Illich BM, Jones AE. Prognosis of emergency department patients with suspected infection and intermediate lactate levels: a systematic review. J Crit Care. 2014;29(3):334–339. doi:10.1016/j.jcrc.2013.12.017.

76. Nelson JL, Harmon ME, Robergs RA. Identifying plasma glycerol concentration associated with urinary glycerol excretion in trained humans. J Anal Toxicol. 2011;35(9):617–623. doi:10.1093/anatox/35.9.617.

77. Röder PV, Wu B, Liu Y, Han W. Pancreatic regulation of glucose homeostasis. Exp Mol Med. 2016;48(3):e219–e219. doi:10.1038/emm.2016.6.

78. Berger C, Zdzieblo D. Glucose transporters in pancreatic islets. Pflugers Arch. 2020;472(9):1249–1272. doi:10.1007/s00424-020-02383-4.

79. Vieira E, Salehi A, Gylfe E. Glucose inhibits glucagon secretion by a direct effect on mouse pancreatic alpha cells. Diabetologia. 2007;50:370–379. doi:10.1007/s00125-006-0511-1.

80. Vergari E, Knudsen JG, Ramracheya R, Salehi A, Zhang Q, Adam J, et al. Insulin inhibits glucagon release by SGLT2-induced stimulation of somatostatin secretion. Nat Commun. 2019;10(1):139. doi:10.1038/s41467-018-08193-8.

81. van Haeften TW, Boonstra E, Veneman TF, Gerich JE, van der Veen EA. Dose-response characteristics for glucose-stimulated insulin release in man and assessment of influence of glucose on arginine-stimulated insulin release. Metabolism. 1990;39(12):1292–1299. doi:10.1016/0026-0495(90)90186-G.

82. Turco G, Brossa C, D’alberto M, Regis G, Segre G, Bianchi E, et al. Kinetic analysis of the response of plasma glucose, insulin, and C-peptide to glucagon injection in normal and diabetic subjects. Diabetes. 1981;30(8):685–693. doi:10.2337/diab.30.8.685.

83. Tu J, Tuch BE. Glucose regulates the maximal velocities of glucokinase and glucose utilization in the immature fetal rat pancreatic islet. Diabetes. 1996;45(8):1068–1075. doi:10.2337/diab.45.8.1068.

84. Saunders PT, Koeslag JH, Wessels JA. Integral rein control in physiology. J Theor Biol. 1998;194(2):163–173. doi:10.1006/jtbi.1998.0746.

85. Mandarino LJ, Bonadonna RC, Mcguinness OP, Halseth AE, Wasserman DH. Regulation of muscle glucose uptake in vivo. Compr Physiol. 2010; p. 803–845. doi:10.1002/cphy.cp070227.

86. Dimitriadis GD, Maratou E, Kountouri A, Board M, Lambadiari V. Regulation of postabsorptive and postprandial glucose metabolism by insulin-dependent and insulin-independent mechanisms: an integrative approach. Nutrients. 2021;13(1):159. doi:10.3390/nu13010159.

87. Geser C, Müller-Hess R, Felber J. The insulin: glucagon ratio and the secretion of growth hormone after intravenous administration of amino acids and carbohydrates in healthy subjects. Transfus Med Hemother. 1973;1(6):483–489. doi:10.1159/000219092.

88. Geary N. Postprandial Suppression of Glucagon Secretion: A Puzzlement. Diabetes. 2017;66(5):1123–1125. doi:10.2337/dbi16-0075.

89. Janah L, Kjeldsen S, Galsgaard KD, Winther-Sørensen M, Stojanovska E, Pedersen J, et al. Glucagon receptor signaling and glucagon resistance. Int J Mol Sci. 2019;20(13):3314. doi:10.3390/ijms20133314.

90. Sylow L, Tokarz VL, Richter EA, Klip A. The many actions of insulin in skeletal muscle, the paramount tissue determining glycemia. Cell Metab. 2021;33(4):758–780. doi:10.1016/j.cmet.2021.03.020.

91. Meng ZX, Gong J, Chen Z, Sun J, Xiao Y, Wang L, et al. Glucose sensing by skeletal myocytes couples nutrient signaling to systemic homeostasis. Mol Cell. 2017;66(3):332–344. doi:10.1016/j.molcel.2017.04.007.

92. Adeva-Andany MM, Funcasta-Calderón R, Fernández-Fernández C, Castro-Quintela E, Carneiro-Freire N. Metabolic effects of glucagon in humans. J Clin Transl Endocrinol. 2019;15:45–53. doi:10.1016/j.jcte.2018.12.005.

93. Snyder WS, Cook M, Nasset E, Karhausen L, Tipton IH. Report of the task group on reference man; 1975.

94. Ho KC, Roessmann U, Straumfjord J, Monroe G. Analysis of brain weight. I. Adult brain weight in relation to sex, race, and age. Arch Pathol Lab Med. 1980;104(12):635–639.

95. Chowdhury B, Sjöström L, Alpsten M, Kostanty J, Kvist H, Löfgren R. A multicompartment body composition technique based on computerized tomography. Int J Obes Relat Metab Disord. 1994;18(4):219–234.

96. Heinemann A, Wischhusen F, Püschel K, Rogiers X. Standard liver volume in the Caucasian population. Liver Transpl Surg. 1999;5(5):366–368. doi:10.1002/lt.500050516.

97. Price PS, Conolly RB, Chaisson CF, Gross EA, Young JS, Mathis ET, et al. Modeling interindividual variation in physiological factors used in PBPK models of humans. Crit Rev Toxicol. 2003;33(5):469–503. doi:10.1080/10408440390242324.

98. Krekels EHJ, Rower JE, Constance JE, Knibbe CAJ, Sherwin CMT. Chapter 8 - Hepatic Drug Metabolism in Pediatric Patients. In: Xie W, editor. Drug Metabolism in Diseases. Boston: Academic Press; 2017. p. 181–206.

99. Brundin T, Wahren J. Whole body and splanchnic oxygen consumption and blood flow after oral ingestion of fructose or glucose. Am J Physiol Endocrinol Metab. 1993;264(4):E504–E513. doi:10.1152/ajpendo.1993.264.4.E504.

100. Prinz H. Hill coefficients, dose–response curves and allosteric mechanisms. J Chem Biol. 2010;3:37–44. doi:10.1007/s12154-009-0029-3.

101. Kolb H, Stumvoll M, Kramer W, Kempf K, Martin S. Insulin translates unfavourable lifestyle into obesity. BMC Med. 2018;16:1–10. doi:10.1186/s12916-018-1225-1.

102. Kolb H, Kempf K, Röhling M, Martin S. Insulin: too much of a good thing is bad. BMC Med. 2020;18:1–12. doi:10.1186/s12916-020-01688-6.

103. Li C, Ford ES, McGuire LC, Mokdad AH, Little RR, Reaven GM. Trends in hyperinsulinemia among nondiabetic adults in the US. Diabetes Care. 2006;29(11):2396–2402. doi:10.2337/dc06-0289.

